# Molecular Matchmakers: How ATP and Small Amphiphilic Molecules Fine-Tune FET Proteins Clusters

**DOI:** 10.1101/2025.04.16.649119

**Authors:** Mrityunjoy Kar

**Affiliations:** Leibniz-Institut für Polymerforschung Dresden e.V., Dresden, Germany

**Keywords:** FET proteins, Sub-saturation mesoscale clusters, Biomolecular condensates, ATP modulation, Protein self-assembly, Sequence-specific interactions

## Abstract

FET (FUS-EWSR1-TAF15) family proteins inherently form mesoscale molecular assemblies, known as clusters, under physiological conditions at concentrations well below the threshold for phase separation. This study demonstrates that adenosine triphosphate (ATP), an amphiphilic molecule and essential cellular metabolite, modulates the size of these sub-saturation mesoscale clusters in a concentration-dependent manner. At low concentrations (1-2 mM), ATP acts as a crosslinker for FET proteins, resulting in larger size clusters. At moderate concentrations (5 mM), the size of the clusters decreases but stabilizes. At high concentrations (10 mM), the cluster size further diminishes. Other amphiphilic molecules, including common hydrotropes like sodium xylene sulfonate, sodium toluene sulfonate, and hexanediol, exhibit comparable concentration-dependent effects on FET protein clustering. Notably, these effects cannot be explained solely by hydrotropic or kosmotropic mechanisms; instead, they stem from non-specific interactions between proteins and small molecules. The intrinsic chemical properties of the amphiphilic molecules and FET proteins play a crucial role in regulating mesoscale cluster formation at sub-saturation concentrations.

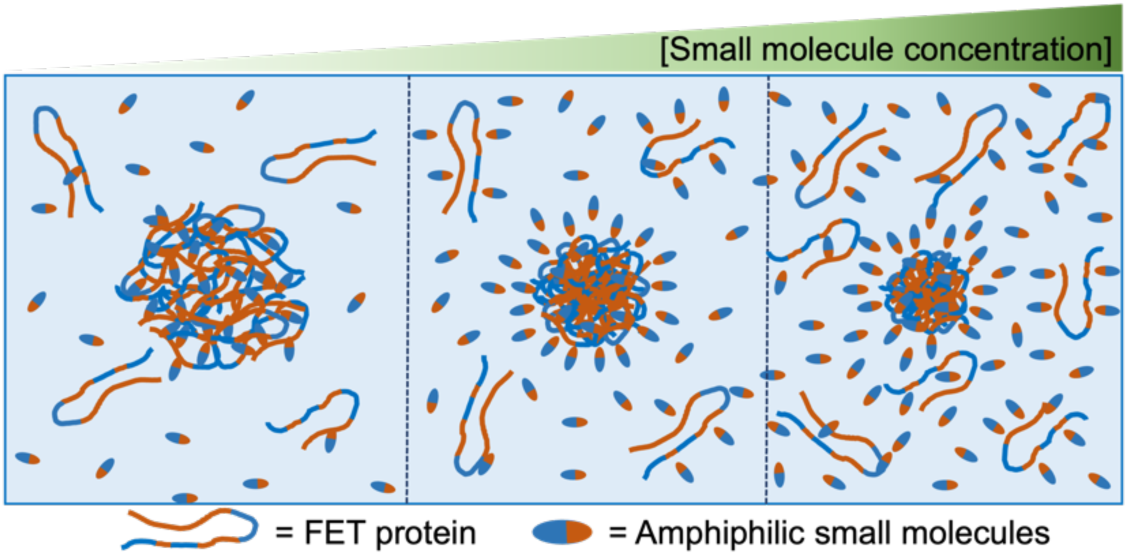

## Introduction

FET proteins are classified as RNA binding proteins, which comprise Fused in Sarcoma (FUS), Ewing Sarcoma RNA binding protein 1 (EWSR1), and TATA-Box Binding Protein Associated Factor 15 (TAF15) ^1^. The FET proteins, present in every cell in our body, exhibit nucleocytoplasmic shuttling and play pivotal roles in RNA biogenesis, such as transcription, splicing, stability, transport, and metabolism ^1^. Members of this protein family demonstrate phase separation phenomena, forming liquid-like states *in vivo* and *in vitro* ^2, 3^, influencing various biological processes, from DNA damage responses to cytoplasmic stress reactions ^4, 5, 6^. Dysregulated phase transitions of FET proteins have been associated with neurodegenerative disorders ^7^. FET proteins are present in cytoplasmic inclusions characteristic of Frontotemporal lobar degeneration (FTLD) and Amyotrophic lateral sclerosis (ALS) ^8^.

FET proteins spontaneously organize into diverse mesoscale molecular assemblies known as clusters, even at concentrations well below the saturation threshold for phase separation ^9, 10^. Similar behavior of sub-saturation cluster formation has been observed in other phase-separating proteins ^11, 12, 13, 14^. Recent investigations have unveiled the prevalence of biomacromolecular mesoscale structures within the cytoplasm ^15^. Specifically, Li and colleagues have highlighted the functional significance of gas vesicle protein clusters in sub-saturated solutions, which impact bacterial fitness ^13^. Given the relevance of sub-saturation mesoscale clusters or condensation in several phase-separating proteins, comprehending the impact of the most abundant cellular metabolite, ATP, on the distribution of sub-saturation mesoscale clusters of FET proteins is imperative.

Adenosine triphosphate, ATP, is an energy currency of biology ^16, 17^. The concentration of ATP in cells stays very high, at an average of 4.4 mM ^18^. ATP consists of adenine (a nitrogenous base), ribose (a five-carbon sugar), and three serially bonded phosphate groups ^16, 17^. ATP releases energy via hydrolysis when needed by converting it into adenosine diphosphate (ADP) and inorganic phosphate^16^. Beyond its role in providing energy, ATP is believed to regulate between soluble and aggregated protein states through its hydrotrope property ^19, 20, 21, 22, 23, 24, 25, 26^.

Hydrotropes are small, amphiphilic molecules that significantly enhance the solubility of sparingly soluble hydrophobic compounds in aqueous solutions ^27, 28^. Hydrotropes typically comprise hydrophilic (water-attracting) and hydrophobic (water-repelling) segments. They have a lower degree of hydrophobicity compared to surfactants and cannot form well-organized structures like micelles. However, hydrotropes can organize around hydrophobic molecules via non-specific interactions and significantly increase the solubility of hydrophobic small molecules in water ^29^. Given the amphiphilic properties, ATP is regarded as a ‘biological hydrotrope,’ which helps maintain the solubility of high-concentration proteins, prevents the formation of, as well as dissolves previously formed protein aggregates ^19^. Moreover, the high cellular ATP levels regulate cellular proteostasis and affect the cytoplasm’s physicochemical properties, such as viscosity, macromolecular crowding, and liquid-liquid phase separation ^30, 31, 32^. Herein, we asked the following question: What is the impact of various concentrations of ATP on sub-saturated FET protein clusters?

In addition to ATP, this study also examines the influence of other small amphiphilic molecules (Figure 1) on forming sub-saturation clusters of FET proteins. We investigate two common hydrotropes, sodium xylene sulfonate (NaXS) and sodium toluene sulfonate (NaTS) (Figure 1), on the formation of sub-saturation clusters. Further, we also investigate another amphiphilic molecule, 1,6-hexanediol (Figure 1), frequently utilized *in vivo* and *in vitro* phase separation studies ^33, 34, 35, 36^. Moreover, we demonstrated that a controlled amount of HD and ATP prevents phase separation of FET proteins at saturation concentration and maintains clusters ^9^. Do hydrotropes and HD also influence the formation of FET protein sub-saturation clusters? If so, is there any similarity in the behavior of these molecules compared to ATP? Using dynamic light scattering (DLS) and nanoparticle tracking analysis (NTA), we investigated the formation of sub-saturation mesoscale clusters with and without ATP and other small molecules. All the data were systematically compared to understand the general principles of how the inherent chemistry of both proteins and small molecules influences the formation of mesoscale molecular assemblies (clusters) at various concentrations. Furthermore, we employed Nano differential scanning fluorimetry (NanoDSF) to examine the temperature-induced molecular stability of the protein in the presence of ATP and other small molecules.

**Figure 1:**
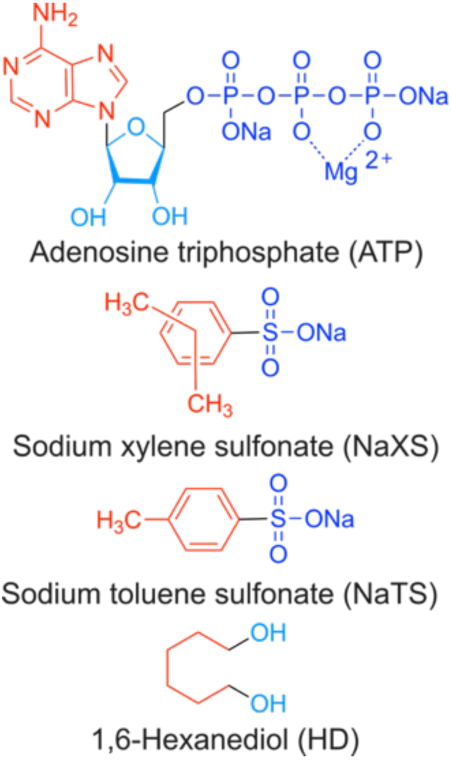
Amphiphilic small molecules: Chemical structure drawing of various molecules, including adenosine triphosphate (ATP), sodium xylene sulfonate (NaXS), sodium toluene sulfonate (NaTS), and hexanediol (HD). In the drawing, the red represents the hydrophobic segments, while the blue represents the hydrophilic segments. Additionally, navy blue depicts more hydrophilic ionic moieties than hydroxy groups, shown in light blue. The NaXS is a mixture of two isomers, where two methyl groups are arranged in either ortho, meta, or para position in the benzene ring.

## Results

### ATP modulates the cluster size distribution of sub-saturation FUS-SNAP clusters

To investigate how ATP and other small amphiphilic molecules influence the formation of mesoscale FET protein clusters at sub-saturation concentrations, we used FUS fused with a C-terminal SNAP-tag (FUS-SNAP). The SNAP-tag increases the saturation concentration of FUS compared to its untagged form, allowing for a broader concentration range to be studied ^3, 9^. We conducted dynamic light scattering (DLS) experiments to monitor sub-saturation cluster formation by measuring the hydrodynamic diameter over 32 minutes, following previous studies ^9, 10^. Figures 2a–c show the hydrodynamic radius over time for FUS-SNAP at 0.25 μM, 0.5 μM, and 1 μM, respectively, in the presence of 10 mM KCl and 20 mM HEPES buffer (pH 7.4). Given our focus on the effects of small amphiphilic molecules on cluster formation, we maintained a low KCl concentration (10 mM), as higher KCl levels inhibit FET protein clustering ^10^.

**Figure 2:**
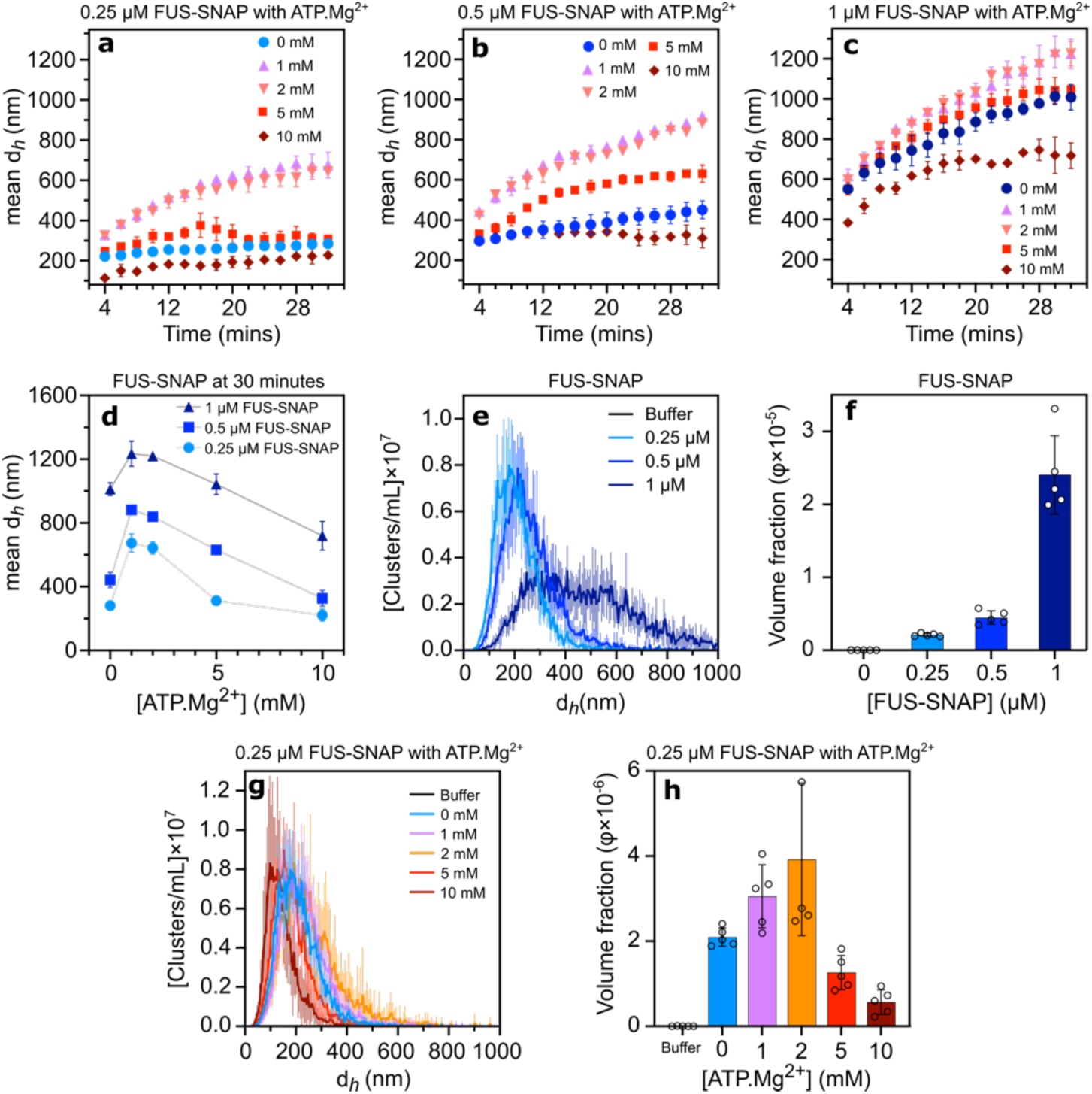
ATP.Mg²⁺ Modulates the sub-saturation cluster size of FUS-SNAP. Dynamic Light Scattering (DLS) data depict the mean hydrodynamic diameter (d*_h_*) of mesoscale clusters at different sub-saturation concentrations of FUS-SNAP (0.25 μM (a), 0.5 μM (b), and 1 μM (c)) over 32 minutes with varying concentrations of ATP.Mg²⁺. The mean hydrodynamic diameter (d*_h_*) of 0.25 μM, 0.5 μM, and 1 μM FUS-SNAP at 30 minutes with varying ATP.Mg²⁺ concentrations (d). Nanoparticle Tracking Analysis (NTA) shows d*_h_* of mesoscale clusters at 0.25 μM, 0.5 μM, and 1 μM FUS-SNAP (e) in 10 mM KCl and 20 mM HEPES pH 7.4. The volume fraction of mesoscale clusters is shown for different FUS-SNAP concentrations (f). NTA data for cluster’ d*_h_* at 0.25 μM FUS-SNAP with different ATP.Mg²⁺ concentrations are shown in (g), while (h) displays the corresponding volume fraction of clusters. NTA data are recorded at around 6 to 8 minutes from the time of sample preparation. Data are presented as mean ± SD, with n=3 (DLS) and n=5 (NTA) independent samples.

Our DLS data revealed a time-dependent increase in the mean hydrodynamic diameter of FUS-SNAP mesoscale clusters across all concentrations, with higher FUS-SNAP concentrations leading to a faster rate of cluster growth (Figures 2a–c). Autocorrelation data (Supporting Information 1) confirmed cluster size increases over time without slow modes indicative of cluster-cluster coalescence at low salt concentrations. However, at 1 μM FUS-SNAP, the cluster size exceeded one micron over time, aligning with phase separation under these conditions, unlike the sub-saturation clusters observed at 0.25 μM and 0.5 μM. The average hydrodynamic diameters of FUS-SNAP clusters at 30 minutes were approximately 281 nm, 441 nm, and 1021 nm for 0.25 μM, 0.5 μM, and 1 μM FUS-SNAP, respectively (Figure 2d).

To complement the DLS findings, we performed nanoparticle tracking analysis (NTA), which provides both size distributions and particle concentrations. NTA results were comparable with DLS data, showing smaller cluster sizes at 0.25 μM and 0.5 μM FUS-SNAP compared to 1 μM (Figure 2e, Supporting Table 1). Notably, DLS and NTA data do not overlap for the same sample; NTA data revealed smaller scatterer populations than DLS data ^37^. Volume fraction measurements at 0.2, 0.4, and 2.4 for 0.25, 0.5, and 1 μM FUS-SNAP, respectively (Figure 2f), further indicated cluster formation at lower concentrations and phase separation at 1 μM in 10 mM KCl and 20 mM HEPES (pH 7.4).

For ATP-dependent studies, we used ATP·Mg²⁺, the biologically active complex formed by ATP binding to magnesium ions ^38^. At 1 mM and 2 mM ATP·Mg²⁺, FUS-SNAP clusters continued to grow over 32 minutes (Figures 2a–c). Cluster size increased by 1.2- to 2.3-fold in 1 mM ATP·Mg²⁺ over 30 minutes relative to buffer-only conditions (Figure 2f), depending on FUS-SNAP concentration. Notably, no slow modes were detected in the correlation functions (Supporting Information 2, 3, 4), suggesting that low ATP·Mg²⁺ concentrations promoted larger assemblies without cluster-cluster coalescence. In the presence of higher concentrations of ATP·Mg²⁺, the cluster size was reduced compared to 1 and 2 mM ATP·Mg²⁺ in all FUS-SNAP concentrations (Figure 2a-c, f). Notably, at 5 mM and 10 mM ATP·Mg²⁺, cluster growth plateaued around 30 minutes, indicating the stabilization of clusters (Figures 2a–c).

At a sub-saturation concentration of 0.25 μM FUS-SNAP, NTA data mirrored the trends observed in DLS (Figure 2g). Lower ATP·Mg²⁺ concentrations shifted cluster size distributions toward larger sizes, whereas higher ATP·Mg²⁺ concentrations reduced cluster sizes. A similar trend was observed in volume fraction data (Figure 2h), underscoring the role of ATP in modulating mesoscale cluster formation.

As a control, we examined ATP without Mg²⁺ at 0.25 μM FUS-SNAP (Supporting Information 5). The trends remained similar: low ATP concentrations increased cluster size, while high concentrations reduced it relative to buffer-only conditions, though the effect required lower ATP concentrations than ATP·Mg²⁺. At 0.5 mM ATP, cluster size increased, whereas at 1 mM ATP, it decreased compared to buffer-only conditions (Supporting Information 5a). DLS data were inconclusive at 5 and 10 mM ATP conditions as the clustering was significantly inhibited. However, NTA data followed the expected trend: cluster size increased at 0.5 mM ATP and decreased at higher concentrations (Supporting Information 5b). At lower ATP concentrations, volume fraction data aligned well with cluster size, showing an increase at 0.5 mM ATP and a decrease at 1 mM ATP compared to buffer-only conditions (Supporting Information 5c). At 5 and 10 mM ATP, clusters of a larger size formed with lower populations, leading to an overall increase in volume fraction (Supporting Information 5b).

### Regulation of sub-saturation FUS-SNAP clusters by ADP and AMP but not adenosine

ATP consists of three phosphate groups linked to the 5’ carbon of the sugar molecule of adenosine (Figure 1). To pinpoint the key moieties responsible for cluster regulation at 0.25 μM FUS-SNAP, we progressively removed phosphate groups from ATP and tested adenosine diphosphate (ADP), adenosine monophosphate (AMP), and adenosine (Figure 3a). We observed trends similar to ATP.Mg²⁺ in the presence of ADP at equivalent concentrations (Figure 3b, e). At 1 mM AMP, cluster sizes increased compared to the buffer-only condition. However, AMP significantly inhibited cluster formation at higher concentrations (5 and 10 mM), producing obscure DLS data (Figure 3c, e). In contrast, adenosine-induced only slight size increases at all concentrations compared to the buffer-only condition, with no discernible concentration-dependent trend (Figure 3d, e).

**Figure 3:**
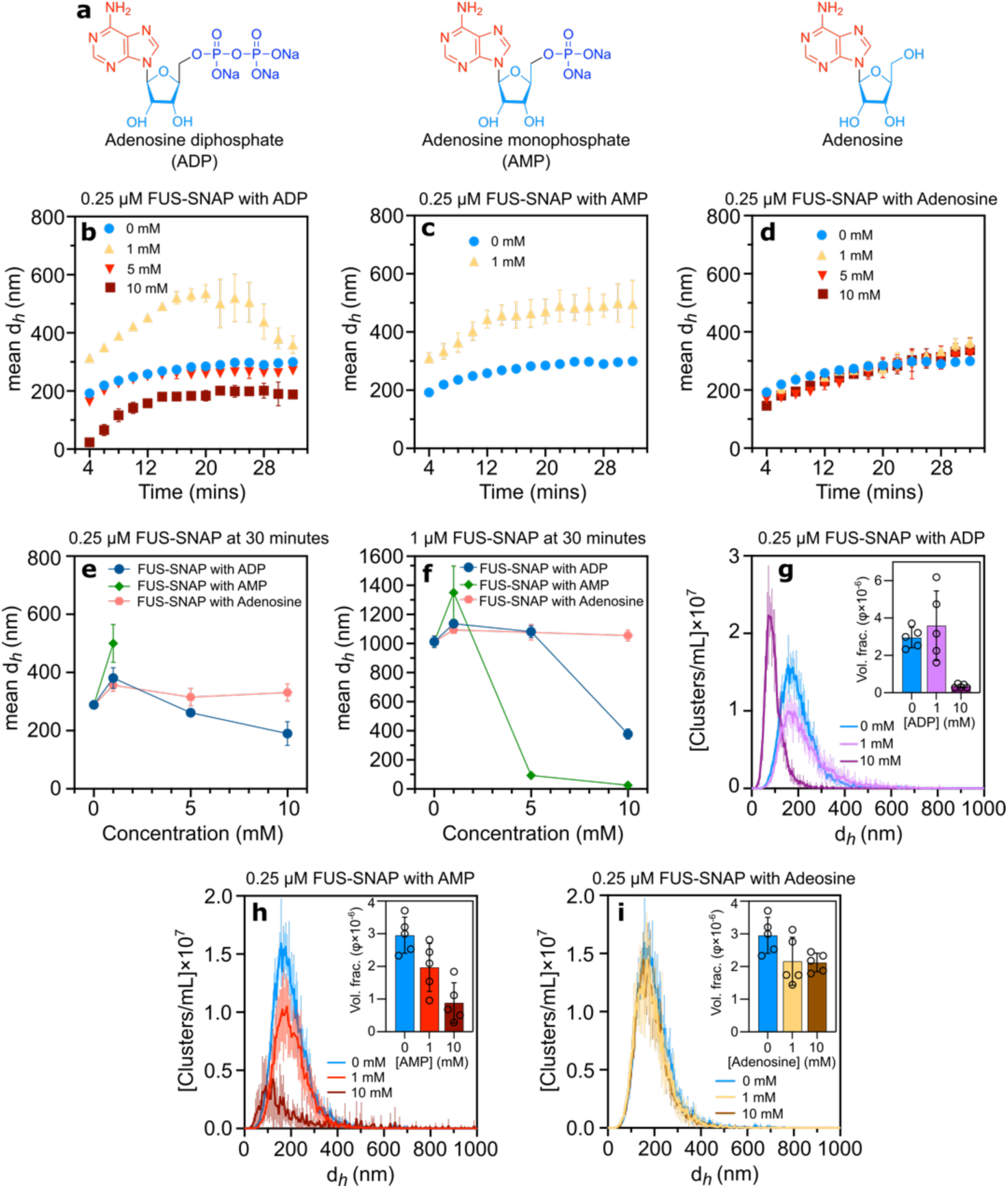
Adenosine diphosphate (ADP) and adenosine monophosphate (AMP) modulate the size of sub-saturation FUS-SNAP clusters, whereas adenosine shows no regulatory effect. Chemical structures of ADP, AMP, and adenosine (a). Dynamic Light Scattering (DLS) data showing mean hydrodynamic diameters (d*_h_*) of mesoscale clusters at 0.25 μM FUS-SNAP over 32 minutes in the presence of various concentrations of ADP (b), AMP (c), and adenosine (d). DLS data shows a mean d*_h_* of 0.25 μM FUS-SNAP (e) and 1 μM FUS-SNAP (f) at 30-minute time points with different concentrations of ADP, AMP, and adenosine. Nanoparticle Tracking Analysis (NTA) data at 0.25 μM FUS-SNAP for clusters’ d*_h_* with ADP (g), AMP (h), and adenosine (i); insets show corresponding volume fractions of the cluster. NTA data are recorded at around 6 to 8 minutes from the time of sample preparation. Data are presented as mean ± SD from n = 3 (DLS) and n = 5 (NTA) independent experiments.

Further investigation at 1 µM FUS-SNAP revealed similar trends: cluster sizes gradually increased over time in the presence of low concentrations of ADP and AMP compared to buffer-only conditions and vice versa (Supporting Information 6). In the case of adenosine, 1 µM FUS-SNAP cluster size increased slightly at equivalent concentrations of ATP.Mg²⁺, no consistent size trends were observed (Supporting Information 6). As shown in Figure 3f, cluster sizes increased at low concentrations but decreased with increasing concentrations of ADP and AMP (5 and 10 mM) compared to the 1 mM condition. Similar to the 0.25 µM, at 1 µM FUS-SNAP, the presence of adenosine also exhibits similar trends: the condensate size increases across all concentrations compared to the buffer-only condition, without a discernible concentration-dependent pattern (Figure 3f).

NTA data corroborated the DLS findings at 0.25 µM FUS-SNAP, revealing consistent trends. At 1 mM ADP, cluster size distributions showed a slight increase, whereas at 10 mM ADP, the distributions decreased compared to buffer-only conditions (Figure 3g). The volume fraction followed a similar pattern, increasing at 1 mM ADP and decreasing at 10 mM ADP (Figure 3g). At 1 mM AMP, the cluster size distribution was slightly larger (Figure 3h), though the volume fraction of clusters was lower than in the buffer-only condition (Figure 3h). At 10 mM AMP, the cluster size distribution and volume fraction significantly decreased compared to buffer-only conditions (Figure 3h). In the presence of adenosine, no clear trends were observed in cluster size distributions at different concentrations (Figure 3i). Additionally, the volume fractions at 1 and 10 mM adenosine remained similar but were reduced compared to buffer-only conditions. These observations underscore the essential role of phosphate moieties in regulating sub-saturation FUS-SNAP cluster formation.

### The phosphate moiety in nucleotides plays a pivotal role in regulating the size of sub-saturated FUS-SNAP clusters

To investigate how the phosphate group in ATP, ADP, and AMP influences cluster size, we examined the effects of sodium tripolyphosphate (STPP), sodium pyrophosphate (SPP), and sodium phosphate (SP) (Figure 4a) on the sub-saturation cluster formation of FUS-SNAP. Notably, at low STPP, SPP, and SP concentrations, the hydrodynamic diameter of the 0.25 µM FUS-SNAP sub-saturation clusters increased over time, mirroring the effects of ATP.Mg²⁺, ADP, and AMP when compared to buffer-only conditions (Figures 4b-d, e). However, at higher concentrations (5 and 10 mM), the cluster size of 0.25 µM FUS-SNAP decreased and stabilized over time, in contrast to the buffer-only condition (Figures 4b-d, e). The formation of clusters at 1 µM FUS-SNAP in the presence of STPP, SPP, and SP displayed similar trends to those observed at sub-saturation concentrations (Supporting Information 7, Figure f).

**Figure 4:**
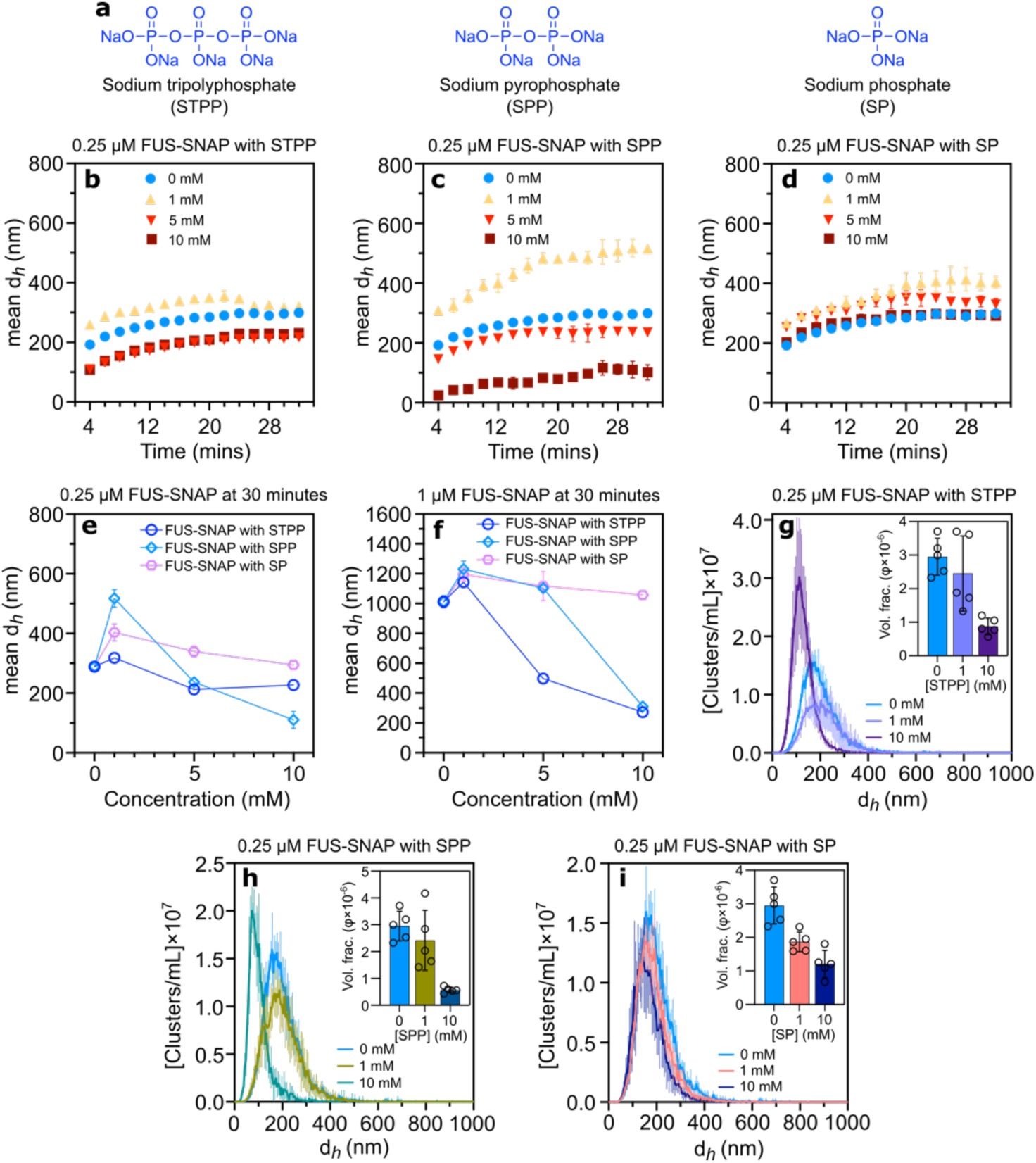
Phosphates are salient in controlling the size of sub-saturated FUS-SNAP clusters. Chemical structure of sodium tripolyphosphate (STPP), sodium pyrophosphate (SPP), and sodium phosphate (SP) (a). Dynamic Light Scattering (DLS) data show mesoscale clusters’ mean hydrodynamic diameter (d*_h_*) at 0.25 mM FUS-SNAP over 32 minutes with various concentrations of sodium tripolyphosphate (STPP) (b), sodium pyrophosphate (SPP) (c), and sodium phosphate (SP) (d). DLS data shows a mean d*_h_* of 0.25 mM (e) and 1 mM (f) FUS-SNAP at 30-minute time points with various STPP, SPP, and SP concentrations. Nanoparticle Tracking Analysis (NTA) data at 0.25 μM FUS-SNAP for clusters’ d*_h_* with STPP (g), SPP (h), and SP (i); insets show corresponding volume fraction of clusters. NTA data are recorded at around 6 to 8 minutes from the time of sample preparation. Data are presented as mean ± SD from n = 3 (DLS) and n = 5 (NTA) independent experiments.

Furthermore, Figure 4g illustrates the NTA data showing size distributions of 0.25 µM FUS-SNAP in the presence of 1 and 10 mM STPP. At 1 mM STPP, the size distribution shifted toward larger clusters, whereas at 10 mM STPP, the distribution narrowed, and the volume fraction of clusters decreased compared to the buffer-only condition. A similar pattern was observed with 0.25 µM FUS-SNAP in the presence of SPP (Figure 4h). In contrast, with SP, the cluster size distributions remained relatively consistent across all concentrations, though increasing SP concentrations led to a decrease in the volume fraction of clusters (Figure 4i). This data suggests that the phosphate moieties (STPP, SPP, and SP) regulate FUS-SNAP cluster formation in a concentration-dependent manner, with distinct effects on cluster size and volume fraction.

### Similar to ATP.Mg²⁺, common hydrotropes, also regulate the cluster size of sub-saturated FUS-SNAP clusters

ATP is classified as a ‘biological hydrotrope’ due to its behavior resembling typical hydrotropes ^19^. To investigate this further, we examined the effects of the common hydrotropes sodium xylene sulfonate (NaXS) and sodium toluene sulfonate (NaTS) on sub-saturation cluster formation. As illustrated in Figure 5a, the hydrodynamic diameters of 0.25 µM and 1 µM FUS-SNAP clusters in the presence of NaXS and NaTS exhibit trends similar to ATP.Mg²⁺. At lower concentrations, these hydrotropes promote an increase in cluster size, while at higher concentrations, the cluster size decreases compared to buffer-only conditions (Supporting Information 7). However, the concentration ranges required for NaXS and NaTS to produce these effects are significantly higher than those required for ATP.Mg²⁺. At lower FUS-SNAP concentrations (0.25 µM), cluster inhibition was observed at 40 mM NaXS and NaTS (Figure 5a, Supporting Information 8). For 1 µM FUS-SNAP, inhibition occurred at around 60 mM (Figure 5a, Supporting Information 8).

**Figure 5:**
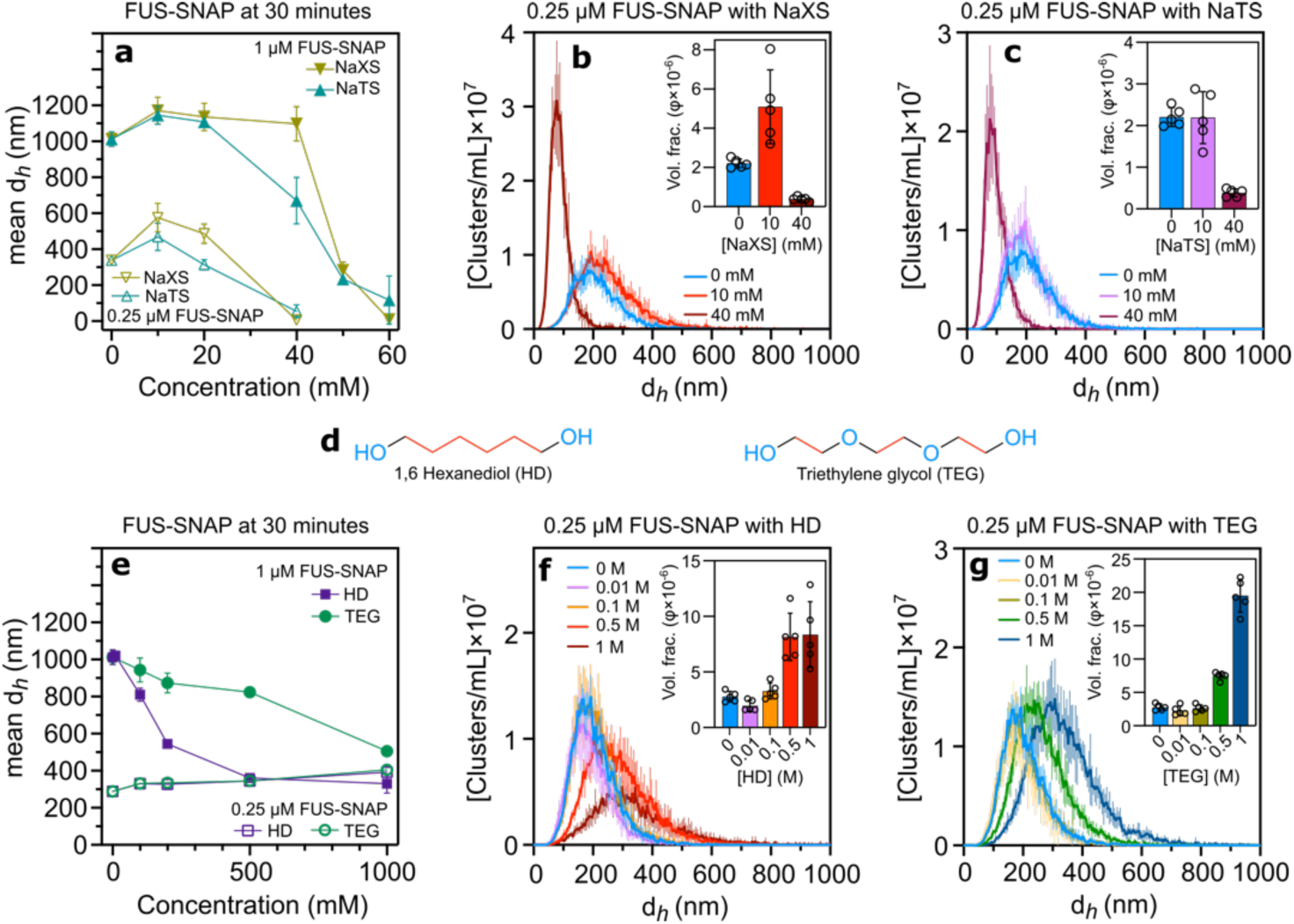
Common hydrotropes—Sodium Xylene Sulfonate (NaXS), Sodium Toluene Sulfonate (NaTS), Hexanediol (HD), and Triethylene Glycol (TEG)—also influence the size of sub-saturation clusters. Dynamic Light Scattering (DLS) data show mesoscale clusters’ mean hydrodynamic diameter (*d_h_*) of 0.25 μM and 1 μM FUS-SNAP at 30-minute time points with various concentrations of Sodium Xylene sulfonate (NaXS) and Sodium Toluene Sulfonate (NaTS) (a). Nanoparticle Tracking Analysis (NTA) data at 0.25 μM FUS-SNAP for clusters’ d*_h_* with NaXS (b) and NaTS (c); insets show corresponding volume fraction of clusters. Chemical structure of HD and TEG (d). DLS data show mesoscale clusters’ mean d*_h_* of 0.25 μM and 1 μM FUS-SNAP at 30-minute time points with various concentrations of HD and TEG (e). NTA data at 0.25 μM FUS-SNAP for clusters’ d*_h_* with HD (f) and TEG (g); insets show the corresponding volume fraction of cluster. NTA data are recorded at around 6 to 8 minutes from the time of sample preparation. Data are presented as mean ± SD from n = 3 (DLS) and n = 5 (NTA) independent experiments.

Additionally, NTA data correlated well with DLS measurements at 0.25 µM FUS-SNAP. At 10 mM NaXS, the cluster size distribution shifted toward larger sizes, while at 40 mM NaXS, the cluster size decreased significantly compared to the buffer-only condition (Figure 5b). The cluster volume fraction followed a similar pattern: a higher value at 10 mM NaXS and a marked decrease at 40 mM NaXS. NaTS exhibited comparable behavior (Figure 5c) to NaXS. These findings demonstrate that common hydrotropes, like NaXS and NaTS, mimic the cluster-regulating effects of ATP.Mg²⁺ but require higher concentration ranges to exert similar effects.

### Hexanediol regulates the cluster size of FUS-SNAP clusters

We next investigated 1,6-hexanediol (HD) due to its amphiphilic structural similarity to ATP and hydrotropes (Figure 1). HD consists of six linked methylene groups (hydrophobic) with hydroxyl moieties (hydrophilic) at both ends (Figure 5d). To compare its effects, we also tested triethylene glycol (TEG), which has the same number of methylene groups as HD but with oxygen atoms separating each ethylene unit, thereby reducing hydrophobicity (Figure 5d).

Interestingly, HD and TEG exhibited different trends of effects depending on FUS-SNAP concentrations. At sub-saturation concentration (0.25 μM FUS-SNAP), HD and TEG slightly increased cluster size with increasing concentrations compared to the buffer-only condition (Supporting Information 8, Figure 5e). Conversely, at the phase separation concentration (1 μM FUS-SNAP), higher concentrations of HD and TEG led to a significant decrease in cluster size compared to the buffer-only condition (Figure 5e). Notably, HD had a more substantial effect on cluster size reduction than TEG. At 0.5 and 1 M concentrations for both compounds, the cluster size diminished and stabilized compared to lower concentrations and the buffer-only condition (Supporting Information 9).

NTA data corroborated the DLS findings. As shown in Figures 5f and 5g, increasing concentrations of HD and TEG, respectively, caused a shift toward larger cluster size distributions at 0.25 μM FUS-SNAP compared to buffer-only conditions. The volume fraction of clusters followed a similar pattern (Figures 5f, 5g). These results indicate that significantly high concentrations of HD and TEG regulate FUS-SNAP clusters at higher protein concentrations. However, HD and TEG show limited or no effect on lower concentrations of FUS-SNAP clusters.

### Sub-saturated clusters formation of untagged FUS with ATP and hydrotropes exhibit similar trends to FUS-SNAP

To confirm that the SNAP-tag does not alter the clustering behavior of untagged FUS in the presence of small molecules, we examined sub-saturation cluster formation using untagged FUS at varying concentrations of ATP.Mg²⁺ and other small molecules. At 0.25 μM FUS in buffer containing 10 mM KCl and 20 mM HEPES (pH 7.4), clusters formed and progressively increased in mean size over time, similar to FUS-SNAP (Supporting Information 10). Autocorrelation analysis further confirmed this gradual size increase over 32 minutes, indicating extended decay times (Supporting Information 10).

In the presence of 1 mM ATP.Mg²⁺, the hydrodynamic diameter of 0.25 μM FUS progressively increased over time compared to the buffer-only condition (Supporting Information 9). At 5 mM ATP.Mg²⁺, the mean cluster size initially increased compared to the buffer-only condition but subsequently decreased and stabilized over time (Figure 6a). At 7.5 mM ATP.Mg²⁺, the mean cluster size started at a smaller size and gradually increased, eventually stabilizing, resembling the 5 mM condition (Supporting Information 10). At 10 mM ATP.Mg²⁺, DLS measurements were unable to detect cluster formation due to interference from the high ATP.Mg²⁺ concentration.

**Figure 6:**
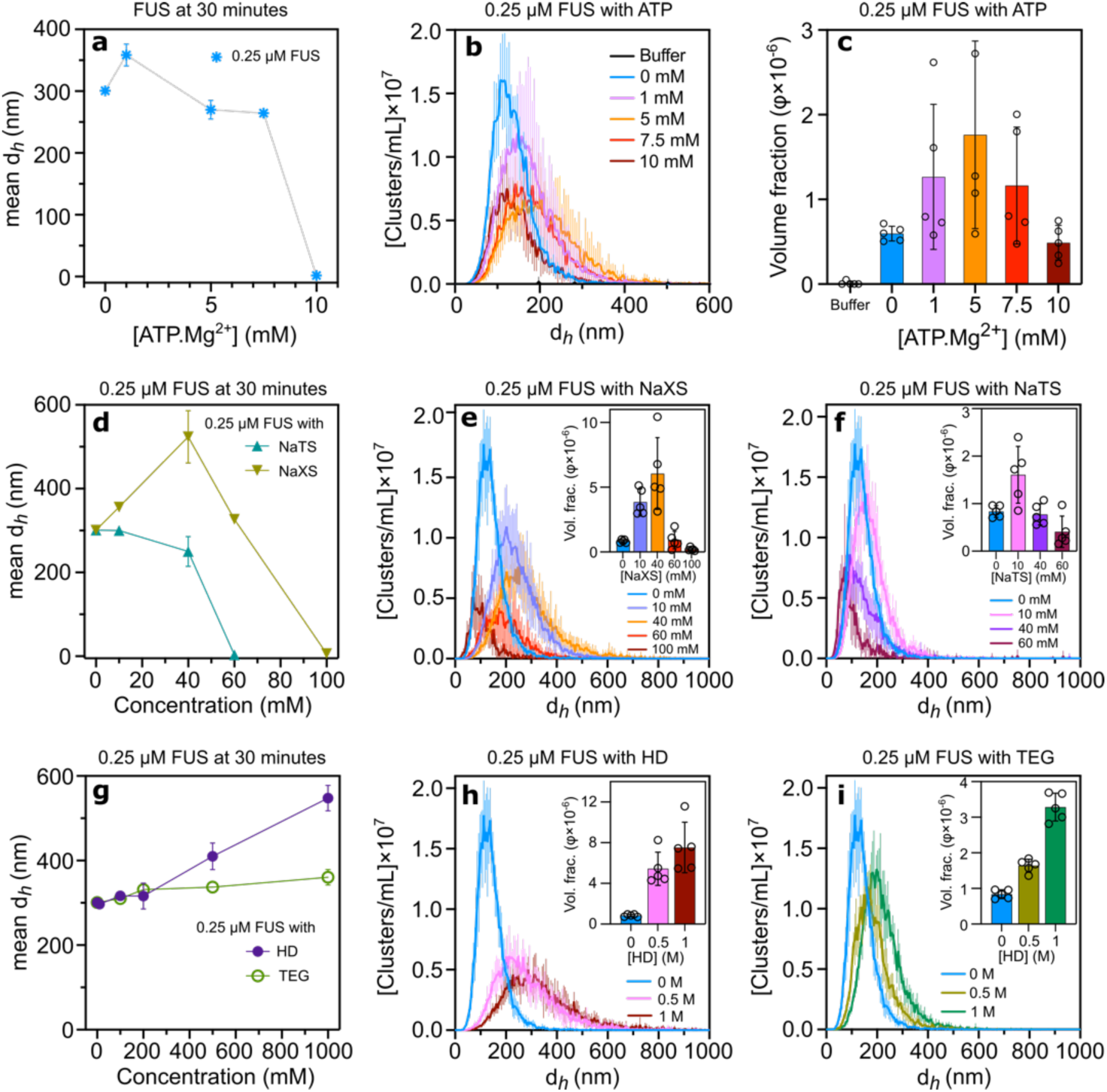
Sub-saturation cluster formation of untagged FUS is similar to FUS-SNAP in the presence of ATP.Mg²⁺, hydrotopes (NaXS and NaTS), HD, and TEG. Dynamic Light Scattering (DLS) data show mesoscale clusters’ mean hydrodynamic diameter (d*_h_*) of 0.25 mM FUS at 30-minute time points with various concentrations of ATP.Mg^2+^ (a). Nanoparticle Tracking Analysis (NTA) data for cluster hydrodynamic diameter at 0.25 μM FUS with different ATP.Mg²⁺ concentrations are shown in (b), while (c) displays the corresponding volume fraction of clusters. DLS data show mesoscale clusters’ mean d*_h_* at 0.25 mM FUS at 30-minute time points with various concentrations of NaTS and NaXS (d). NTA data at 0.25 μM FUS for clusters’ d*_h_* with NaXS (e) and NaTS (f); insets show the corresponding volume fraction of clusters. DLS data show mesoscale clusters’ mean d*_h_* at 0.25 mM FUS at 30-minute time points with various concentrations of HD and TEG (g). NTA data at 0.25 μM FUS for clusters’ d*_h_* with HD (h) and TEG (i); insets show the corresponding volume fraction of clusters. NTA data are recorded at around 6 to 8 minutes from the time of sample preparation. Data are presented as mean ± SD from n = 3 (DLS) and n = 5 (NTA) independent experiments.

The DLS data shown in Figure 6a is consistent with FUS-SNAP observations. Lower ATP.Mg²^+^concentrations promote increased cluster size, while higher concentrations lead to size reduction (Figure 6a). However, the effect of ATP.Mg²+ at various concentrations on FUS is distinctly different from that on FUS-SNAP. NTA data corroborated with DLS findings (Figure 6b). Unlike DLS, at 10 mM ATP.Mg²⁺, smaller clusters were detected in NTA experiments (Figure 6b). Furthermore, the volume fraction of clusters increased at lower ATP.Mg²⁺ concentrations and decreased at higher concentrations (Figure 6c). These findings demonstrate that ATP.Mg²⁺ actively modulates sub-saturation cluster formation in untagged FUS, exhibiting a concentration-dependent pattern that is both similar to and distinct from FUS-SNAP observations.

### Impact of NaXS and NaTS on untagged FUS cluster formation

We next examined how NaXS and NaTS influence the clustering behavior of untagged FUS. As shown in Figure 6d, the clustering trends in the presence of NaXS were similar to those observed with FUS-SNAP. At lower NaXS concentrations, cluster size increased, while higher concentrations led to a decrease in cluster size. In the presence of 10 and 40 mM NaXS, the mean cluster size of 0.25 μM FUS increased compared to the buffer-only condition. However, unlike FUS-SNAP, untagged FUS formed clusters even at 60 mM NaXS, and clustering was inhibited at 100 mM NaXS (Figure 6d, Supporting Information 9). Unlike NaXS, at 10 mM NaTS, cluster size remained comparable to buffer-only conditions. As the NaTS concentration increased, cluster size progressively decreased, and cluster formation was fully inhibited at 60 mM NaTS (Figure 6d, Supporting Information 10). These findings indicate that NaXS and NaTS modulate the clustering of untagged FUS in a concentration-dependent manner, similar to FUS-SNAP, but exhibit subtle differences in clustering thresholds at higher hydrotrope concentrations. NTA data further corroborated these observations. The cluster size distributions and volume fraction of 0.25 μM untagged FUS significantly increased with increasing NaXS concentrations till 40 mM and decreased further, increasing concentration to 100 mM (Figure 6e). As NaTS concentrations increase, the cluster size distributions and volume fraction of NTA data exhibit similar trends, as evidenced by the DLS data (Figures 6d and 6f).

### Effect of HD and TEG on untagged FUS cluster formation

In the presence of 100 mM HD, the mean cluster size of 0.25 μM untagged FUS showed a slight increase compared to buffer-only conditions, similar to the behavior observed with equivalent concentration FUS-SNAP. As the HD concentration increased, the cluster size of untagged FUS grew significantly (Figure 6g, Supporting Information 10). Notably, unlike 0.25 μM FUS-SNAP, untagged FUS exhibited a pronounced increase in cluster size at 500 mM and 1000 mM (1 M) HD (Figure 6g). In contrast, equivalent TEG concentrations induced only a slight increase in the cluster size of 0.25 μM untagged FUS across all tested concentrations, resembling the trend observed with FUS-SNAP (Figure 6g, Supporting Information 9).

NTA data further corroborated these observations. The cluster size distributions and volume fraction of 0.25 μM untagged FUS significantly increased with increasing HD concentrations (Figure 6h). With increasing TEG concentrations, the cluster size distributions and volume fraction also increased compared to buffer-only conditions (Figure 6i). However, in the presence of TEG, the distributions were narrower compared to those observed with HD. These findings suggest that the SNAP-tag, as a ∼20 kDa protein tag, influences clustering behavior by altering sequence-specific interactions with small molecules. While clustering trends in response to ATP, hydrotropes, HD, and TEG were broadly similar for FUS-SNAP and untagged FUS, notable differences were explicitly observed in response to HD, emphasizing the impact of the SNAP-tag on interrelates with small molecules.

### The sequence composition of FET family proteins modulates their response to ATP and other small molecules, influencing sub-saturation clustering

FUS proteins belong to the FET family of RNA-binding proteins, characterized by their intrinsic disorder and RNA-binding domains ^1^. Despite structural similarities, variations in protein sequences and molecular weights confer unique chemical identities to these proteins. To gain further insights, we explored the sub-saturation cluster formation of EWSR1-SNAP and TAF15-SNAP, similar to FUS-SNAP, in the presence of ATP.Mg^2+^, and other small molecules. The SNAP-tagged versions were used for easier purification and handling, as the tag increased the saturation concentration required for phase separation.

### Effect of ATP.Mg²⁺ on EWSR1-SNAP cluster formation

EWSR1-SNAP exhibited cluster formation trends comparable to FUS-SNAP in the presence of ATP.Mg²⁺. At lower ATP.Mg²⁺ concentrations, the cluster size increased, while higher concentrations led to size reduction compared to buffer-only conditions (Figure 7a). At 1 mM ATP.Mg²⁺, the cluster size of 0.3 μM EWSR1-SNAP increased progressively over time (Supporting Information 10). In contrast, at 5 and 10 mM ATP.Mg²⁺, the cluster size initially decreased and then reached a plateau over time, suggesting higher ATP.Mg²⁺ concentrations stabilize sub-saturation clusters (Supporting Information 11). NTA data confirmed these similar trends found in DLS. At 1 mM ATP.Mg²⁺, cluster size distributions increased compared to buffer-only conditions (Figure 7b). However, with 5 and 10 mM ATP.Mg²⁺, cluster size distributions decreased. The volume fraction of clusters of 0.3 μM EWSR1-SNAP increases at 1 mM ATP.Mg²⁺ and decreases with higher concentrations of ATP.Mg²⁺ (Figure 7c).

**Figure 7:**
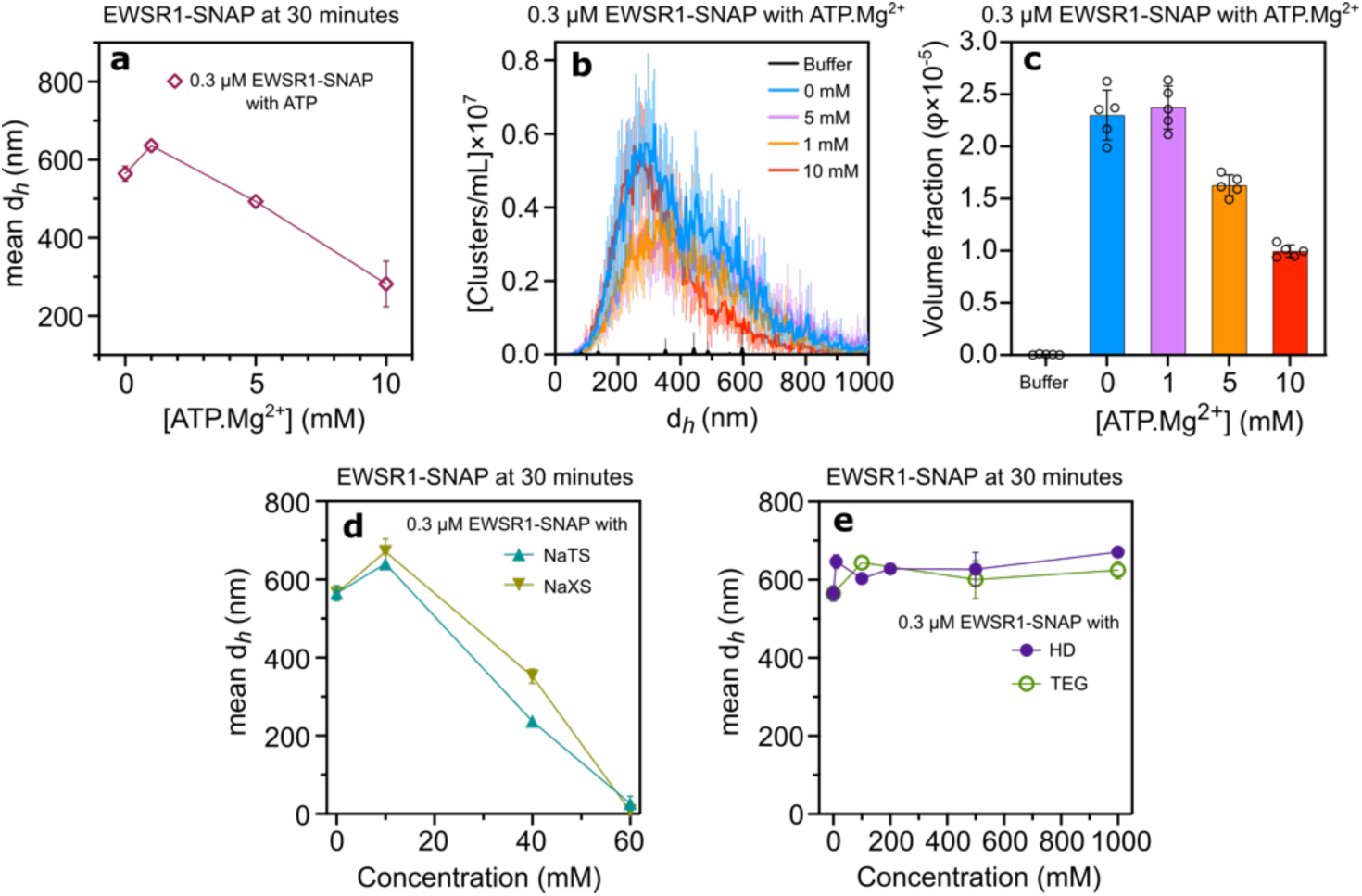
ATP.Mg^2+^, hydrotropes, and HD modulate the size of the Sub-saturation cluster of EWSR1-SNAP. Dynamic Light Scattering (DLS) data show mesoscale clusters’ mean hydrodynamic diameter (*d_h_*) of 0.25 μM EWSR1-SNAP at 30-minute time points with various concentrations of APT.Mg^2+^ (b). Nanoparticle Tracking Analysis (NTA) data for clusters’ d*_h_* at 0.3 μM EWSR1-SNAP with different ATP.Mg²⁺ concentrations are shown in (b), while (c) displays the corresponding volume fraction of clusters. DLS data show mesoscale clusters’ mean d*_h_* at 30-minute time points with various NaTS and NaXS (d), HD, and TEG (e) concentrations. NTA data are recorded at around 6 to 8 minutes from sample preparation time. Data are presented as mean ± SD, with n=3 (DLS) and n=5 (NTA) independent samples.

### Effect of hydrotropes, HD, and TEG on EWSR1-SNAP cluster formation

In the presence of 10 mM NaXS and NaTS, the cluster size of EWSR1-SNAP increased compared to buffer-only conditions (Figure 7d, Supporting Information 10). At 60 mM hydrotrope concentrations, cluster formation was inhibited. Like FUS-SNAP, NaTS had a more substantial effect on cluster inhibition than NaXS, resulting in smaller cluster sizes at equivalent concentrations. At 10 mM HD, the cluster size of EWSR1-SNAP increased compared to the buffer-only condition. However, with higher HD concentrations, the mean cluster size remained similar but larger size compared to the buffer-only condition (Figure 7d, Supporting Information 11). TEG exhibited similar effects to HD, with comparable cluster formation behavior across concentrations (Figure 7e, Supporting Information 10). These findings demonstrate that EWSR1-SNAP follows clustering trends similar to FUS-SNAP in response to ATP.Mg²⁺ and small molecules, with consistent inhibition at high hydrotrope concentrations and stable cluster formation in the presence of HD and TEG.

### Effect of ATP.Mg²⁺ on TAF15-SNAP cluster formation

Similar to FUS-SNAP and EWSR1-SNAP, TAF15-SNAP exhibits comparable trends in sub-saturation cluster formation in the presence of ATP.Mg²⁺. At lower ATP.Mg²⁺ concentrations, the mean cluster size increases relative to buffer-only conditions, whereas higher concentrations lead to a decrease in cluster size (Figure 8a, Supporting Information 12). However, TAF15-SNAP requires higher ATP.Mg²⁺ concentrations to observe these trends compared to FUS-SNAP and EWSR1-SNAP. Notably, at 10 mM ATP.Mg²⁺, the cluster size of 0.25 μM TAF15-SNAP significantly increased over time compared to the buffer-only condition, unlike the stabilization observed with FUS-SNAP and EWSR1-SNAP (Figure 7a). At higher ATP.Mg²⁺ concentrations (15 and 20 mM), cluster sizes decreased and stabilized over time (Supporting Information 12). NTA data corroborated these findings, showing that cluster size distributions initially increased with rising ATP.Mg²⁺ concentrations and then decreased at 20 mM ATP.Mg²⁺ (Figure 8b). The volume fraction of clusters mirrored this trend (Figure 8c).

**Figure 8:**
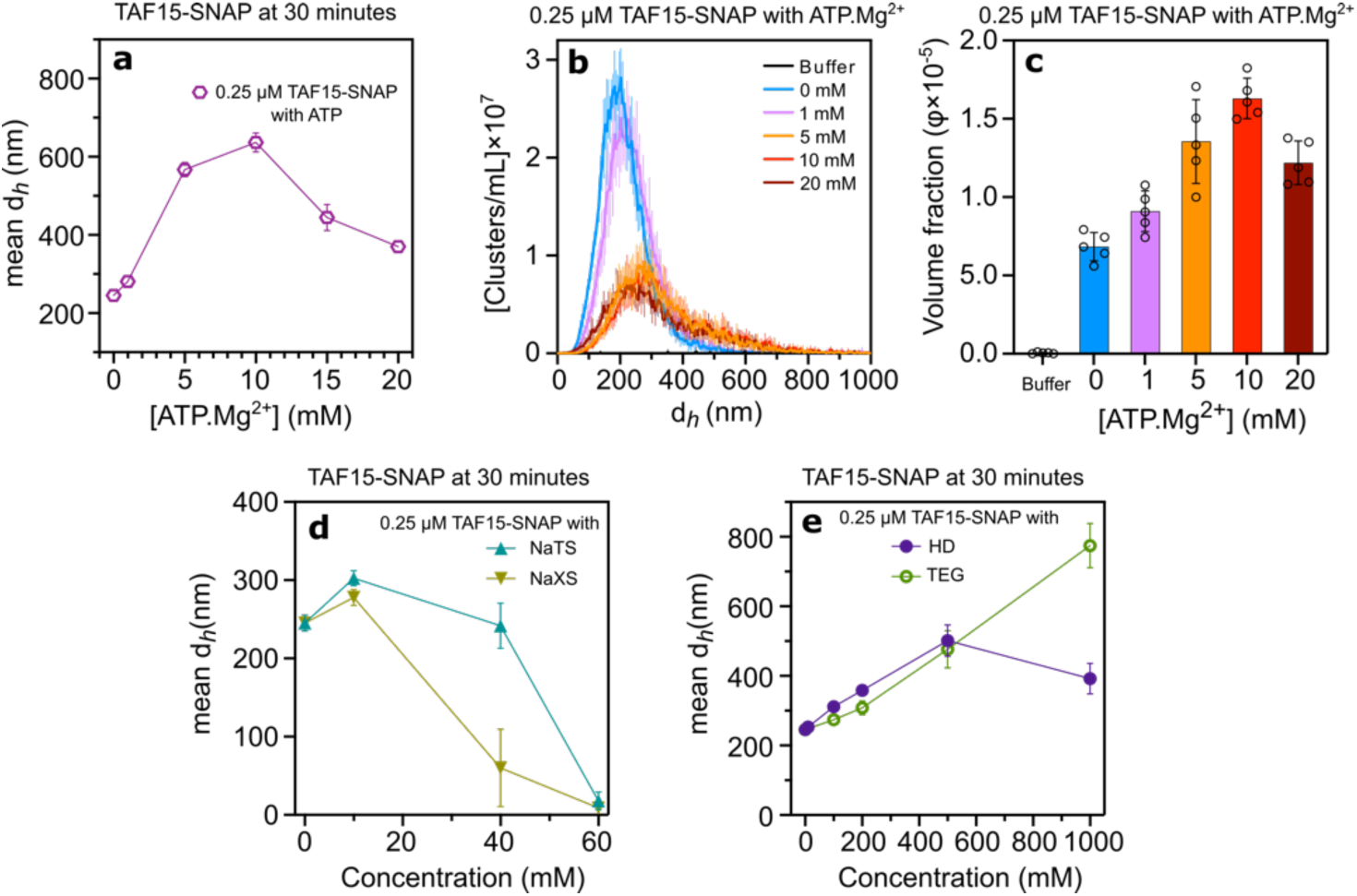
ATP.Mg^2+^, hydrotropes, HD, and TEG modulate the size of the Sub-saturation cluster of TAF15-SNAP. Dynamic Light Scattering (DLS) data show mesoscale clusters’ mean hydrodynamic diameter (*d_h_*) of 0.25 μM TAF15-SNAP at 30-minute time points with various concentrations of APT.Mg^2+^ (b). Nanoparticle Tracking Analysis (NTA) data for clusters’ d*_h_* at 0.25 μM TAF15-SNAP with different ATP.Mg²⁺ concentrations are shown in (b), while (c) displays the corresponding volume fraction of clusters. DLS data show mesoscale clusters’ mean d*_h_* at 30-minute time points with various NaTS and NaXS (d), HD, and TEG (e) concentrations. NTA data are recorded at around 6 to 8 minutes from sample preparation time. Data are presented as mean ± SD, with n=3 (DLS) and n=5 (NTA) independent samples.

### Effect of hydrotropes and HD on TAF15-SNAP cluster formation

In the presence of 10 mM NaXS and NaTS, the TAF15-SNAP cluster size increased compared to buffer-only conditions (Figure 8d, Supporting Information 12). However, as hydrotrope concentrations increased to 40 mM, the cluster size decreased, and at 60 mM, cluster formation was completely inhibited. Interestingly, unlike FUS-SNAP and EWSR1-SNAP, NaXS had a more substantial inhibitory effect on TAF15-SNAP cluster formation than NaTS, resulting in smaller mean cluster sizes.

In the presence of HD, the cluster size of 0.25 μM TAF15-SNAP increased progressively with rising HD concentrations, peaking at 500 mM, where the size was approximately two-fold higher compared to buffer-only conditions (Figure 8e, Supporting Information 12). At 1000 mM (1 M) HD, cluster size decreased compared to the 500 mM condition. Conversely, TEG caused a gradual increase in cluster size with increasing concentrations, albeit less pronounced than HD (Figure 8e, Supporting Information 12).

These findings highlight that sequence-specific chemical properties play a critical role in determining the effects of small molecules on sub-saturation cluster formation in EWSR1-SNAP and TAF15-SNAP. Although common regulatory patterns emerge across FET family proteins, differences in response to ATP.Mg²⁺, hydrotropes, and HD underscore the significance of protein-specific sequence and structural elements.

### Depending on the inherent molecular chemistry ATP.Mg^2+^, NaXS, NaTS, and HD interact differently with FUS-SNAP

ATP is known to prevent heat-induced denaturation of egg white proteins ^19^. Hydrotropes are found to have a concentration-dependent effect on heat coagulation of BSA ^39^. At lower concentrations, hydrotropes stabilize the protein structure, indicative of a higher transition temperature than buffer-only conditions, and at higher concentrations, hydrotropes decrease the transition temperature. Herein, we tested the thermal stability of FUS-SNAP in the presence of ATP, hydrotropes, and HD. We employed nanoscale differential scanning fluorimetry (NanoDSF), a modified method for determining protein stability. This method uses intrinsic tryptophan and tyrosine fluorescence by monitoring two distinct wavelengths, 330 nm, and 350 nm, to analyze the thermal unfolding of the protein. For the NanoDSF experiment, we used 1 μM FUS-SNAP with various concentrations of ATP.Mg^2+^, NaXS, NaTS, and HD.

Our previous report shows that around <1% of proteins are used to form clusters at the sub-saturation concentration below phase separation ^9^. Therefore, the majority of the proteins are monomeric and in solution. For the NanoDSF study, the fraction of proteins in the solution was sufficient to obtain the data represented in Figure 9. At buffer-only conditions, the transition temperature (T_m_) of FUS-SNAP is around 51.9 °C. The temperature-induced denaturation pattern is blueshift, where the fluorescence signal does not change much at 330 nm and changes more strongly at 350 nm (Supporting Information 13). In the presence of 1 mM ATP.Mg^2+^, we observe a similar value of T_m_, around 51.3 °C. At 5 mM ATP.Mg^2+^, the T_m_ increased around 54.2 °C and at 10 mM ATP.Mg^2+^, the T_m_ decreased to around 48.5 °C. In the presence of ATP.Mg^2+^, the denaturation of the proteins is also blueshift (Figure 9a). In the presence of 10 mM NaXS, we observed the transition temperature increase to around 63 °C with a redshift unfolding, where the 330 nm fluorescence intensity changes quite drastically compared to 350 nm (Supporting Information 13). At 60 mM NaXS, we do not observe any transition temperature, where no ratio (330/350) signal was observed (Figure 9b). In the presence of NaTS, both blue and redshift unfolding were recorded with 49.7 °C and 64.4 °C at 10 mM NaTS and 49.5 °C and 65.4 °C at 60 mM NaTS (Figure 9c). In the presence of HD, with increasing concentration, the T_m_ decreases to 51°C, 50.4 °C, and 49.3 °C at 10, 100, and 200 mM HD, respectively, with a blueshift pattern similar to the buffer-only condition (Figure 9d). These data suggest that the interactions between FUS-SNAP and small amphiphilic molecules differ distinctly due to their inherent chemical properties.

**Figure 9:**
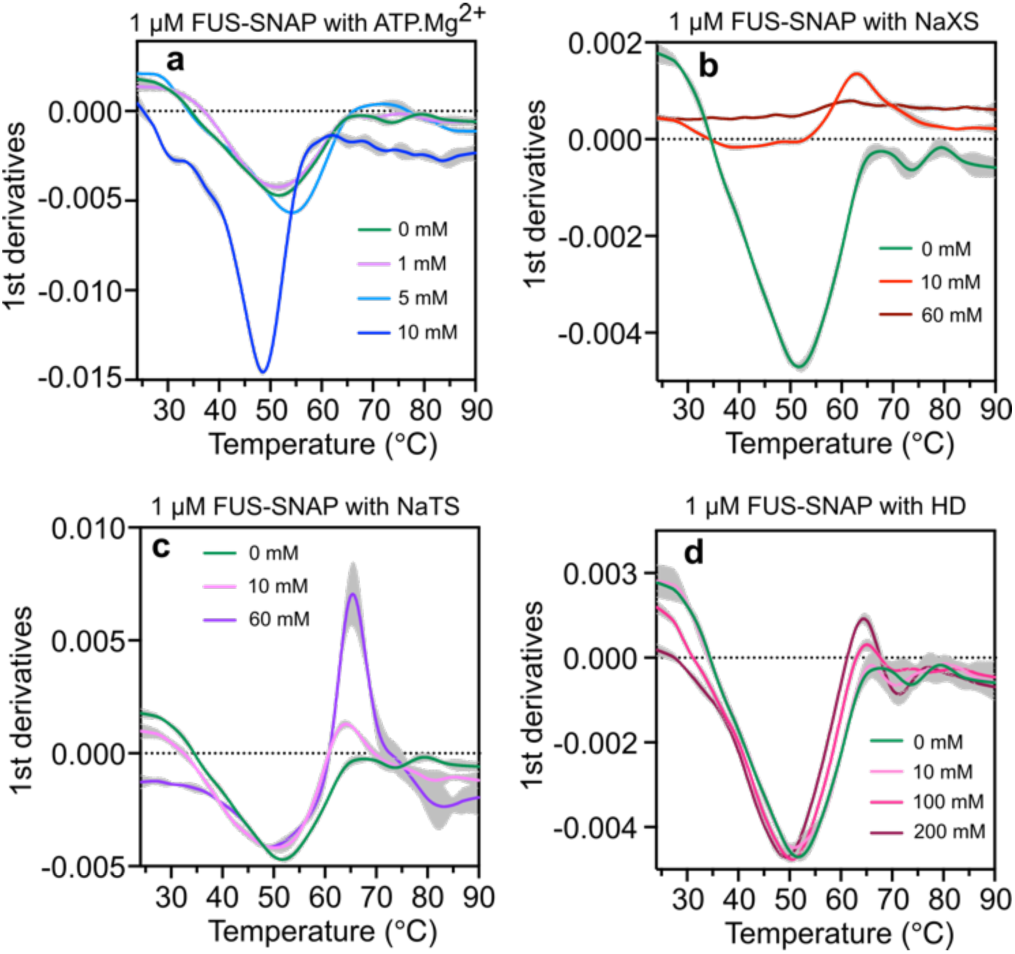
Interactions with ATP and other small amphiphilic molecules alter the transition temperature of FUS-SNAP probed by NanoDSF. 1^st^ Derivatives of the unfolding curves (350/330) ratio with the apparent transition temperatures of FUS-SNAP with ATP.Mg^2+^ (a), NaXS (b), NaTS (c), and HD (d).

## Discussion

Previous studies have shown that ATP.Mg²⁺ dissolves phase-separated droplets of FET proteins at increasing concentrations ^19^. Given its structural and functional similarity to hydrotropes, ATP has been described as a ‘biological hydrotrope.’ ATP also known to enhance liquid-liquid phase separation (LLPS) at lower concentrations (1-2 mM) while dissolving phase-separated condensates at higher concentrations (> 6 mM) ^30^. Additionally, ATP.Mg²⁺ has been found to inhibit the coarsening of phase separation and preserve sub-saturation clusters of FET proteins at saturation concentration ^9^.

In this study, we demonstrated that ATP actively modulates the size of mesoscale clusters in sub-saturation concentrations of FET family proteins. Cluster size increased at lower ATP concentrations but decreased at higher concentrations. This behavior cannot be adequately explained by the hydrotropic effect. Importantly, these mesoscale clusters persisted and stabilized even at physiologically relevant high concentrations of ATP. The enhancement of LLPS at low ATP concentrations was attributed to ATP acting as a bivalent binder through non-specific interactions, where its adenine ring interacts with aromatic residues through π-π or π-cation interactions ^22, 40^, while its triphosphate chain forms electrostatic interactions with arginine or lysine residues ^30^. Similar effects may be attributed to the enhancement of the size of mesoscale clusters at low concentrations of ATP.Mg²⁺ at sub-saturation concentrations of FET proteins.

Furthermore, Mehringer et al. debated that ATP is not a classical hydrotrope but a typical kosmotropic ion (according to the Hofmeister series) ^41^. When comparing the effects of ATP.Mg²⁺, ADP, AMP, STPP, SPP, and SP on cluster formation, we observed similar trends, suggesting these molecules influence sub-saturated mesoscale clusters through analogous mechanisms. Kosmotropic ions are known to interact with water from the hydration layer of proteins, leading to a ‘salting-in’ and ‘salting-out’ mechanism ^41, 42, 43^. Higher concentrations of nucleotides and phosphates inhibit clustering by dissolving or stabilizing the protein. However, the number of phosphate moieties in ATP, STPP, ADP, SPP, AMP, and SP does not correlate well with their effects on sub-saturation clustering. Additionally, at lower concentrations of ATP, STPP, ADP, SPP, AMP, and SP, the kosmotropic effect cannot explain an increase in sub-saturation cluster size. Thus, the Kosmotropic effect alone is insufficient to explain the observed trends. Increasing experimental and computational evidence indicates that kosmotropic ions influence their immediate hydration layers without causing any long-range water ordering ^44^. The Hofmeister series mainly refers to the structure of water in the proximity of ions while leaving out the chemical details of the surfaces of proteins ^44, 45^. So, interactions between FET proteins and small molecules appear critical in modulating cluster size.

Previous studies have highlighted the role of arginine-phosphate interactions in modulating the phase behavior of disordered proteins ^46, 47^. At lower concentrations, arginine-phosphate interactions act as mild crosslinkers, increasing cluster size ^47, 48^. As concentrations increase, these interactions become saturated; therefore, crosslinking effects diminish, and protein monomers are stabilized in solutions, resulting in smaller cluster sizes. The arginine content of FET proteins correlates with these effects: FUS-SNAP (43 arginines), EWSR1-SNAP (53 arginines), and TAF15-SNAP (62 arginines) show differential clustering responses. Higher arginine content in TAF15-SNAP necessitates higher ATP.Mg²⁺ concentrations (15-20 mM) to decrease cluster size.

Hydrotropes such as NaXS and NaTS interact with proteins in a manner similar to ATP·Mg²⁺, but they require concentrations approximately 10 times higher than nucleotides and phosphates to produce comparable effects. This difference arises because proteins have conserved phosphate-binding sites ^49^. Additionally, NaXS and NaTS carry a single negative charge at physiological pH, whereas phosphate has two negative charges, resulting in a higher binding affinity of phosphate for FET proteins compared to NaXS and NaTS. Beyond charge interactions, the hydrophobic xylene and toluene groups in NaXS and NaTS engage in π-π and hydrophobic interactions with the protein. At low concentrations, hydrotropes function as crosslinkers, promoting cluster formation. However, they saturate protein interaction sites at higher concentrations, thereby inhibiting clustering.

The structural differences between NaXS and NaTS influence their regulatory effects. NaXS contains xylene isomers with two methyl groups, making it more hydrophobic than NaTS, which has only one methyl group (toluene). The sequence composition of FUS-SNAP, FUS, EWSR1-SNAP, and TAF15-SNAP varies (Figure 10). While the isoelectric points of FUS-SNAP, FUS, and EWSR1-SNAP are similar, TAF15-SNAP exhibits a distinct isoelectric point. This difference likely contributes to the varying regulatory effects of NaXS and NaTS on FET proteins, highlighting the role of sequence-specific interactions in cluster size regulation.

**Figure 10:**
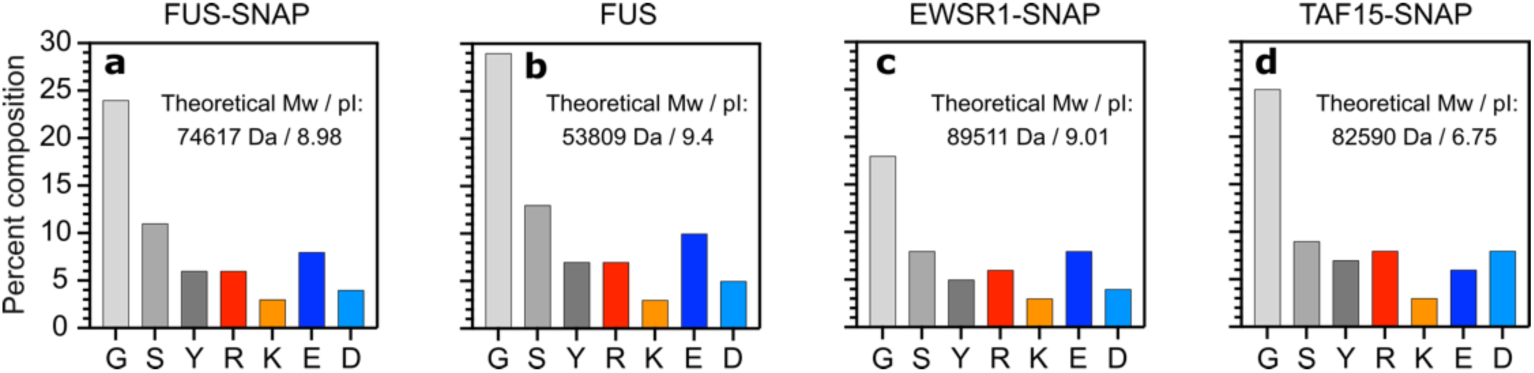
Sequence composition of FET proteins. The percent composition of cationic amino acids—arginine (R) and lysine (K); anionic amino acids—glutamic acid (E) and aspartic acid (D); and the most abundant amino acids—glycine (G), serine (S), and tyrosine (Y) is shown for FUS-SNAP (a), FUS (b), EWSR1-SNAP (c), and TAF15-SNAP (d). The theoretical molecular weight (Mw) and isoelectric point (pI) are reported for each protein.

HD, containing six consecutive hydrophobic methylene groups and weak hydrophilic hydroxy moieties, modulates FET protein clustering and condensate formation by interacting with hydrophobic regions of proteins. This interaction solubilizes and stabilizes the proteins, inhibiting coarsening and condensate formation. At saturation concentrations of FET proteins, increasing HD and TEG concentrations prevent condensation and favor the formation of mesoscale clusters, which is consistent with previous findings ^9^. Despite having the same carbon number as HD, TEG has reduced hydrophobicity due to the presence of three oxygen atoms, resulting in a milder inhibitory effect on condensate formation. At sub-saturation concentrations of FET proteins, both HD and TEG function as mild crosslinkers, increasing cluster size. At very high concentrations (0.5 M and 1 M), HD and TEG also act as molecular crowders, further promoting cluster growth. Notably, sequence differences among FET proteins (Figure 10) modulate their response to HD and TEG, leading to distinctly varied effects. FUS and TAF15-SNAP sub-saturation clusters exhibit a significantly greater size increase compared to FUS-SNAP and EWSR1-SNAP at equivalent concentrations of HD and TEG, highlighting the role of protein sequence-specific chemistry in cluster formation.

NanoDSF analysis revealed that small molecules modulate the denaturation transition temperature (T_m_) of FUS-SNAP. In the buffer-only condition, blueshift denaturation was observed, possibly due to surface-exposed tryptophan or interior tyrosine residues. ATP.Mg²⁺ increased FUS-SNAP stability at 5 mM but decreased stability at 10 mM. NaXS induced a redshift pattern with a 10°C increase in T_m_, indicating strong molecular interactions. NaTS showed dual transitions at 49°C and 64°C, reflecting both blueshift and redshift patterns. HD-induced blueshift patterns resembled those of ATP.Mg²⁺, suggesting similar interaction mechanisms. So, different inherent molecular properties modulate the non-specific interactions with FUS-SNAP differently, thus resulting in different T_m_.

Our findings suggest three concentration-dependent scenarios for sub-saturation cluster modulation:

1. **Low Concentrations:** ATP (1-2 mM), hydrotropes (∼10 mM), and HD (∼100 mM) act as mild crosslinkers, enhancing cluster size compared to buffer-only conditions.
2. **Moderate Concentrations:** Physiological ATP (2-7.5 mM), moderate hydrotrope levels (20-50 mM), and high HD concentrations (0.2-1 M) stabilize cluster sizes by plausibly arranging amphiphilic molecules to inhibit further cluster growth. In earlier studies, we reported that the clusters are more hydrophobic than soluble proteins based on Bis-ANS studies ^10^. At moderate concentrations, these small amphiphilic molecules plausibly arrange themselves to stabilize more hydrophobic mesoscale clusters and inhibit their further growth over time.
3. **High Concentrations:** Excess ATP and hydrotropes saturate protein interaction sites, inhibiting cluster formation to some extent. However, smaller size and lower volume fractions of mesoscale clusters of FET proteins persist at very high concentrations of ATP and hydrotropes.

The non-monotonic phase behavior upon increasing the concentration of an additional component shows similarities with the behavior observed in polymers in two-component solvent systems with a cosolvent that preferentially interacts with polymers/proteins ^50, 51, 52^. Here, an increase in cosolvent concentration induces polymer condensation, followed by re-entry behavior at higher cosolvent levels. Simulation results align with predictions from the adsorption-attraction mean-field theory, which rationalizes and quantitatively predicts the observed behavior ^53, 54^. These studies support a collective transition driven by weak, transient monomer-cosolvent-monomer bridging, which can be explained using mean-field concepts. This phase segregation mechanism, influenced by nonspecific attractive interactions between polymers and smaller components, was previously designated as *gluonic* ^55^. The findings of the present study suggest that the biological hydrotrope ATP, common hydrotropes such as NaXS and NaTS, along with HD and TEG, function as gluonic components, playing a role in the condensation of FET proteins.

## Conclusion

In cellular environments, ATP concentrations vary across tissues, ranging from 2.88 mM in the brain to 7.47 mM in cardiac muscle ^18^. These variations suggest that ATP may differentially modulate FET protein clustering depending on the cell type. Our study demonstrates that ATP and small amphiphilic molecules regulate the size of FET protein clusters in a concentration-dependent manner. At low concentrations, these molecules act as mild crosslinkers, enhancing cluster formation. At intermediate concentrations, they stabilize cluster sizes, while at high concentrations, they saturate protein interaction sites, leading to partial dissolution of clusters while still preserving mesoscale assemblies. These findings challenge conventional hydrotropic and kosmotropic models, highlighting the significance of sequence-specific interactions between FET proteins and small molecules in mesoscale cluster regulation. The structural and chemical properties of both proteins and small molecules influence clustering, emphasizing the importance of inherent molecular interactions in condensate formation. Notably, ATP and hydrotropes function as gluonic components, mediating the condensation and stability of FET protein clusters in a manner reminiscent of polymer behavior in two-component solvent systems. By revealing a nuanced molecular mechanism for protein clustering, this study provides crucial insights into the physicochemical regulation of biomolecular condensates. These findings have broad implications for cellular homeostasis, protein assembly dynamics, and metabolic regulation. Understanding how small amphiphilic molecules modulate cluster formation in both similar and distinct ways opens avenues for selectively regulating mesoscale clusters, potentially offering novel strategies for controlling protein assemblies in biological and synthetic systems. Moreover, FET proteins are also responsible for pathological aggregations in neurodegenerative diseases, such as FTLD and ALS ^8^. The lower ATP concentration in the brain may predispose FET proteins to aggregation, potentially contributing to disease onset. Thus, our findings offer a framework for understanding both physiological and pathological protein assembly, paving the way for targeted strategies to regulate mesoscale clusters in biological and synthetic systems.

## Associated Content

### Supporting Information

The Supporting Information includes detailed materials and methods, supplementary tables, and additional figures supporting the main manuscript’s findings.

## Acknowledgments

I sincerely thank Anthony A. Hyman, Jens-Uwe Sommer, Albena Lederer, Carsten Werner, Susanne Boye, Tyler Harmon, and Jens Rieger for their valuable discussions and insights. I am also grateful to the Hyman Laboratory at the Max Planck Institute for Molecular Cell Biology and Genetics (MPI-CBG) for providing the DNA constructs for all proteins, as well as the PEPC facility at MPI-CBG for their support with protein expression, purification, and characterization.

## Supporting Information

### Materials

**Supporting Table 1:**
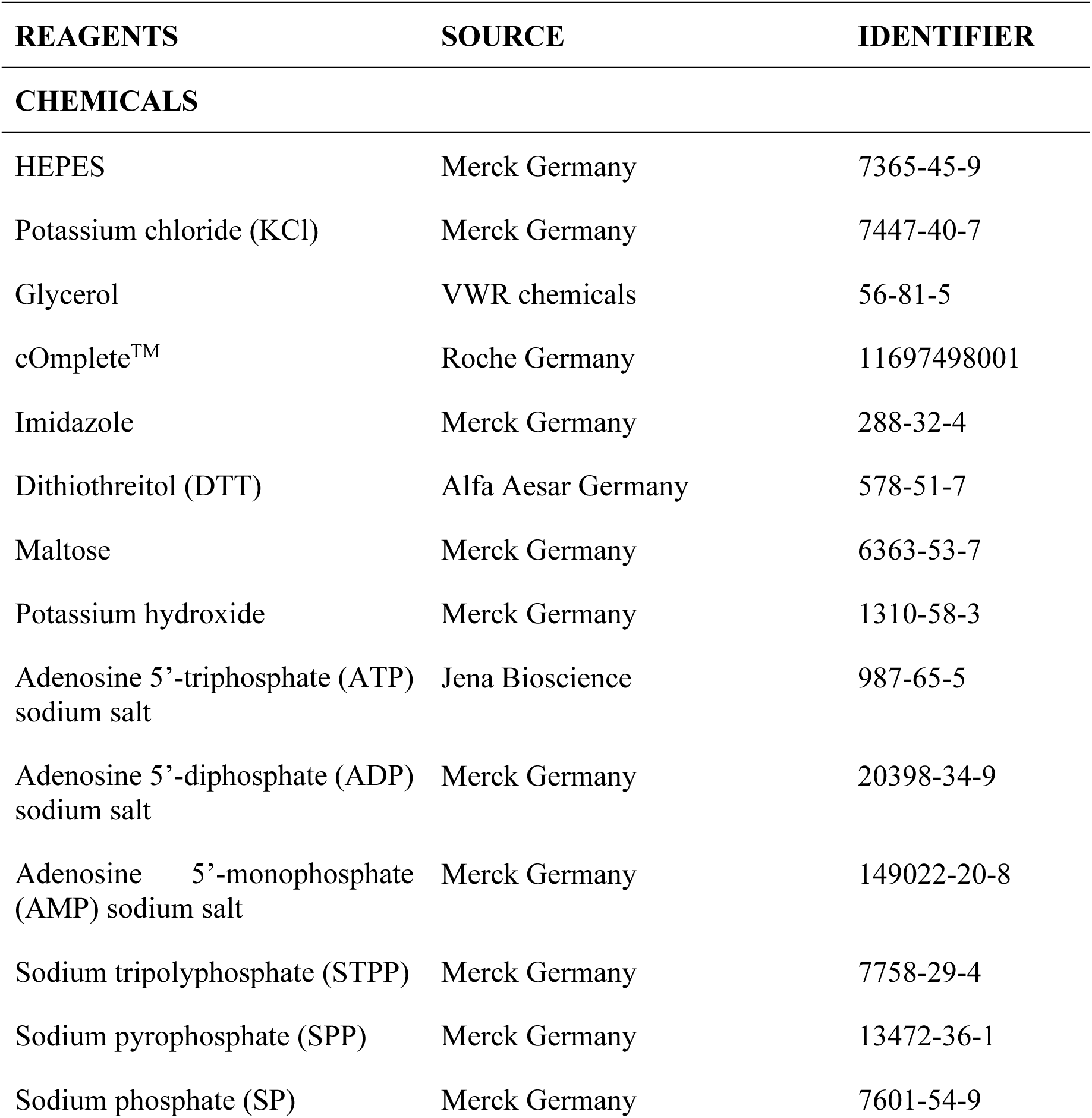
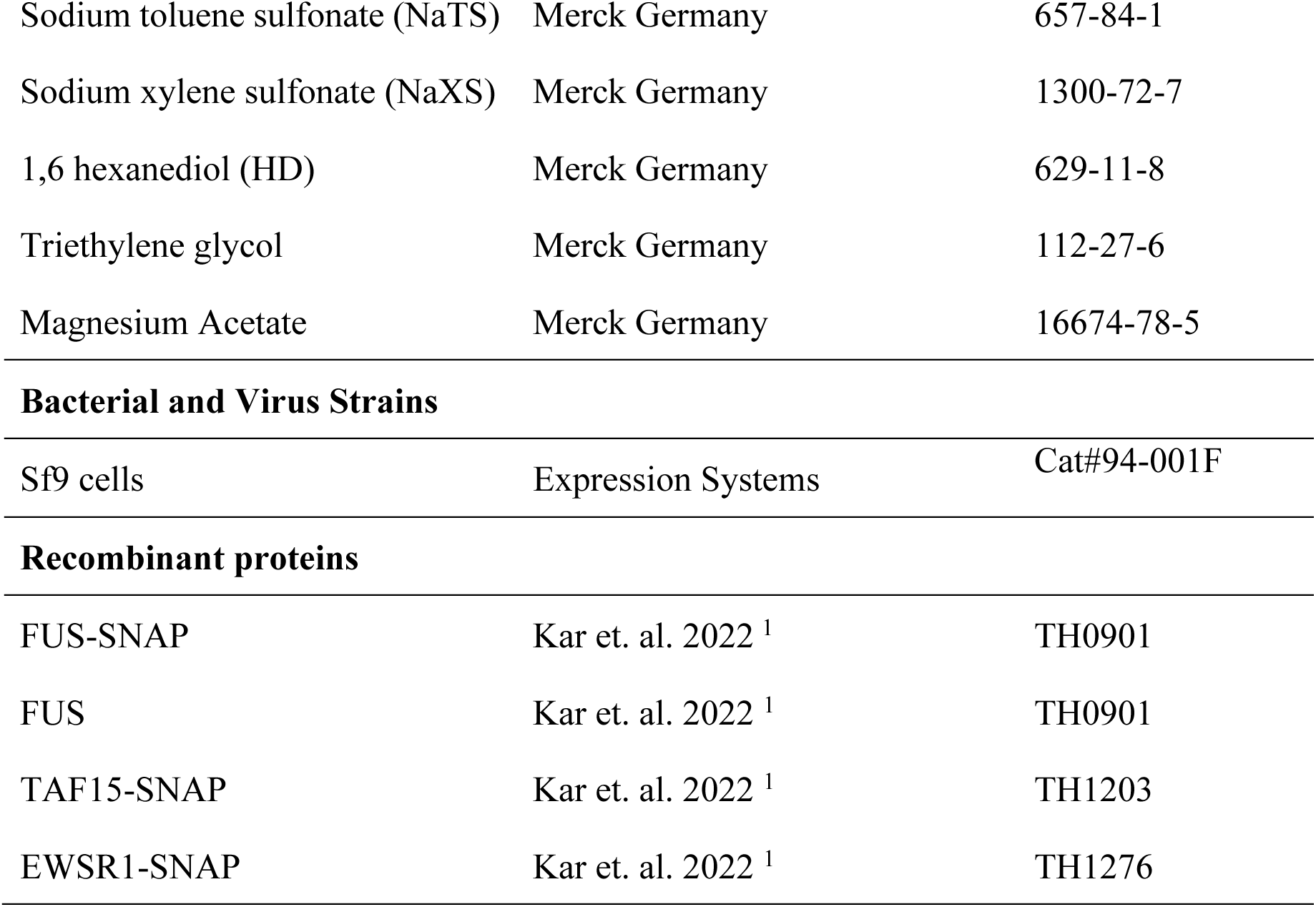
List of reagents, sources, and vendor identifiers, if any.

| REAGENTS | SOURCE | IDENTIFIER |
| --- | --- | --- |
| **CHEMICALS** |  |  |
| HEPES | Merck Germany | 7365-45-9 |
| Potassium chloride (KCl) | Merck Germany | 7447-40-7 |
| Glycerol | VWR chemicals | 56-81-5 |
| cOmplete™ | Roche Germany | 11697498001 |
| Imidazole | Merck Germany | 288-32-4 |
| Dithiothreitol (DTT) | Alfa Aesar Germany | 578-51-7 |
| Maltose | Merck Germany | 6363-53-7 |
| Potassium hydroxide | Merck Germany | 1310-58-3 |
| Adenosine 5'-triphosphate (ATP) sodium salt | Jena Bioscience | 987-65-5 |
| Adenosine 5'-diphosphate (ADP) sodium salt | Merck Germany | 20398-34-9 |
| Adenosine 5'-monophosphate (AMP) sodium salt | Merck Germany | 149022-20-8 |
| Sodium tripolyphosphate (STPP) | Merck Germany | 7758-29-4 |
| Sodium pyrophosphate (SPP) | Merck Germany | 13472-36-1 |
| Sodium phosphate (SP) | Merck Germany | 7601-54-9 |
| Sodium toluene sulfonate (NaTS) | Merck Germany | 657-84-1 |
| Sodium xylene sulfonate (NaXS) | Merck Germany | 1300-72-7 |
| 1,6 hexanediol (HD) | Merck Germany | 629-11-8 |
| Triethylene glycol | Merck Germany | 112-27-6 |
| Magnesium Acetate | Merck Germany | 16674-78-5 |
| **Bacterial and Virus Strains** |  |  |
| Sf9 cells | Expression Systems | Cat#94-001F |
| **Recombinant proteins** |  |  |
| FUS-SNAP | Kar et. al. 2022 <sup>1</sup> | TH0901 |
| FUS | Kar et. al. 2022 <sup>1</sup> | TH0901 |
| TAF15-SNAP | Kar et. al. 2022 <sup>1</sup> | TH1203 |
| EWSR1-SNAP | Kar et. al. 2022 <sup>1</sup> | TH1276 |

#### Amino acid sequences of proteins used in studies

##### 1. FUS-SNAP: This sequence includes full-length FUS (unshaded), a linker that is cleavable by a TEV protease (shaded in yellow), and the SNAP (shaded in gray)

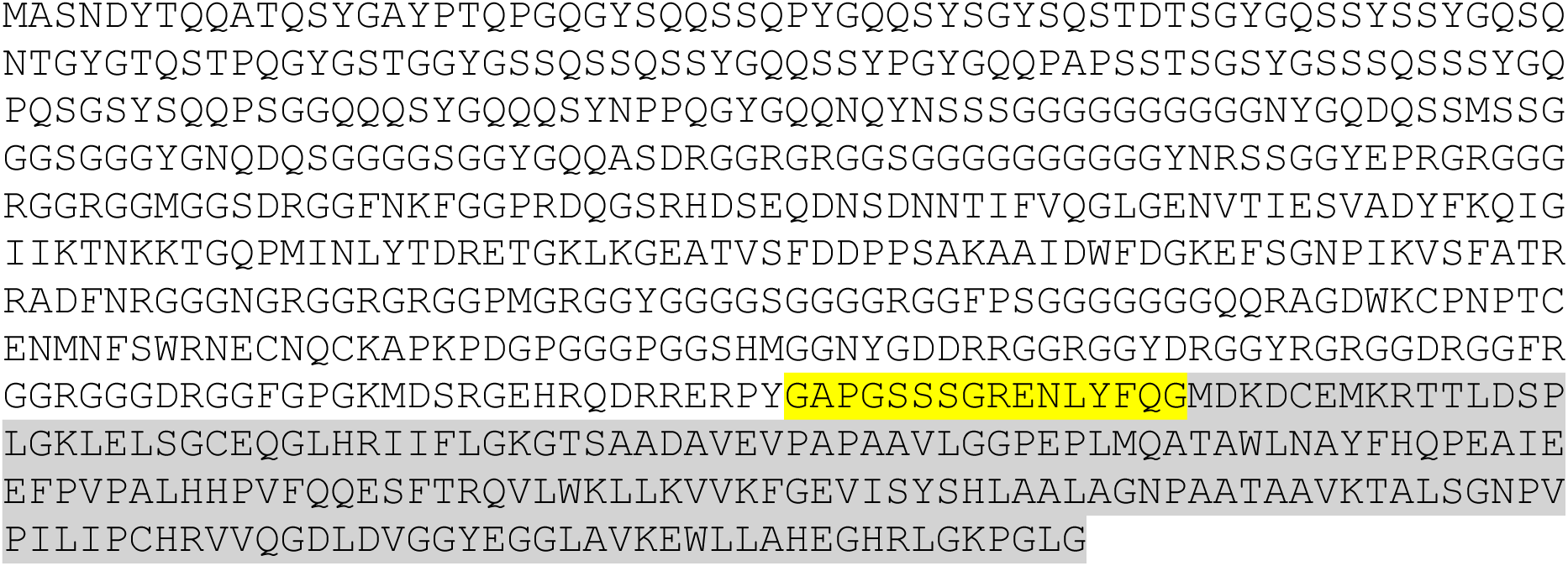

##### 2. FUS

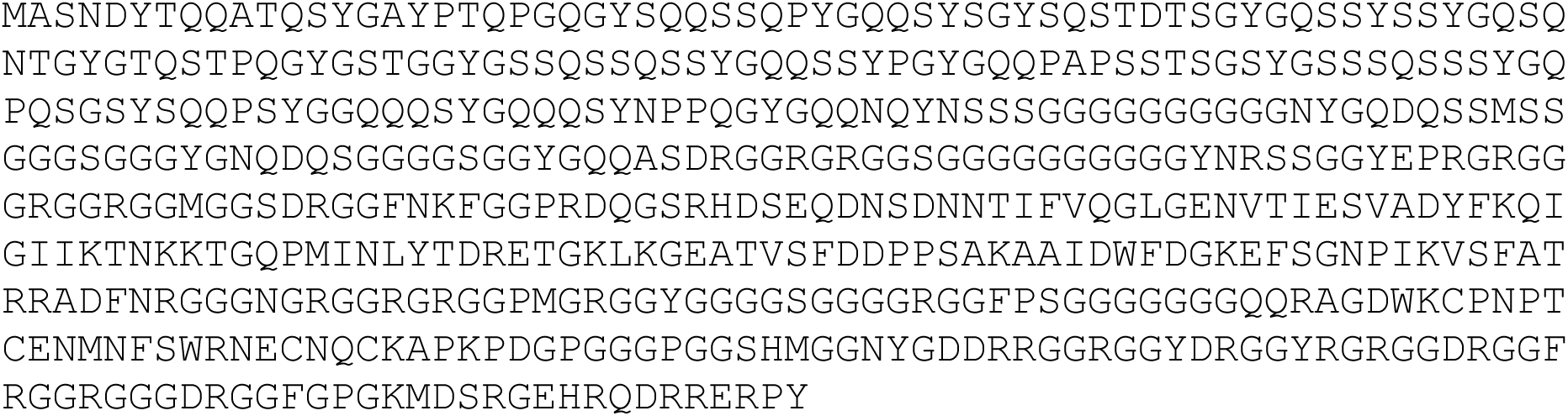

##### 3. TAF15-SNAP: This sequence includes full-length Taf15 (unshaded), a linker that is cleavable by a TEV protease (shaded in yellow), and the SNAP tag (shaded in gray)

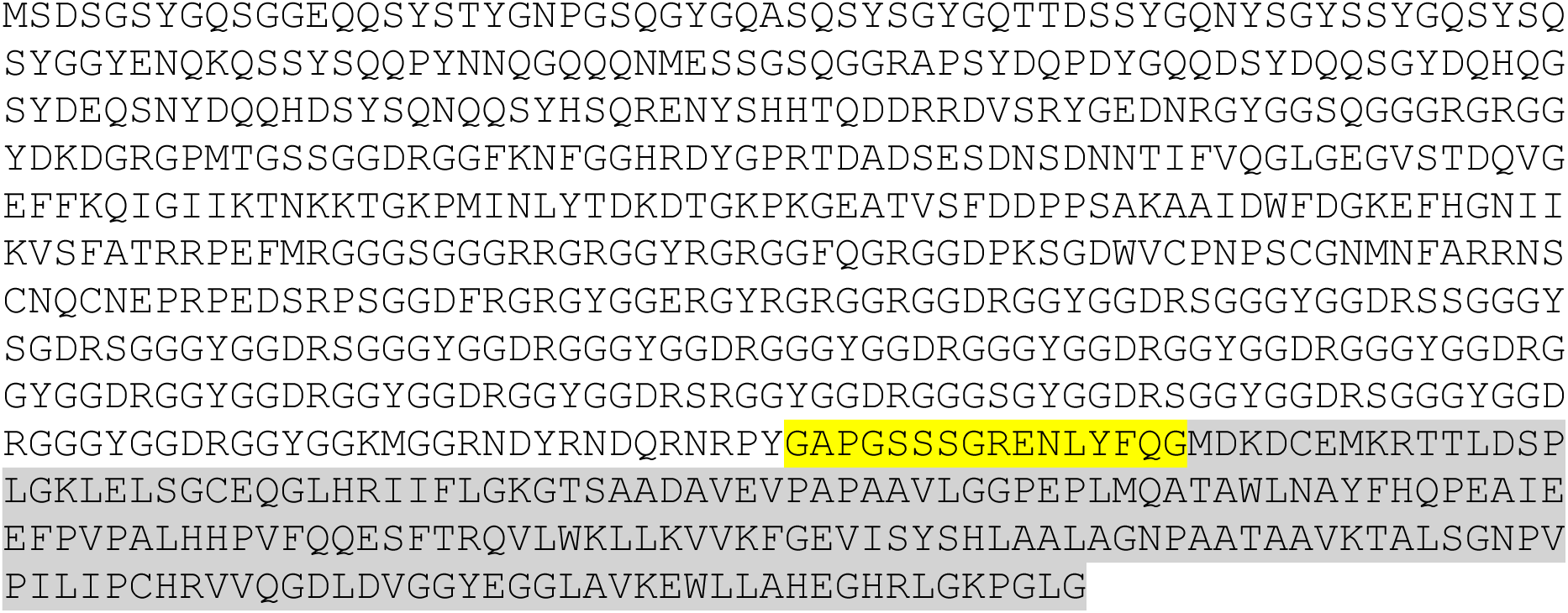

##### 4. EWSR1-SNAP: This sequence includes full-length Ewsr1 (unshaded), a linker that is cleavable by a TEV protease (shaded in yellow), and the SNAP-tag (shaded in gray)

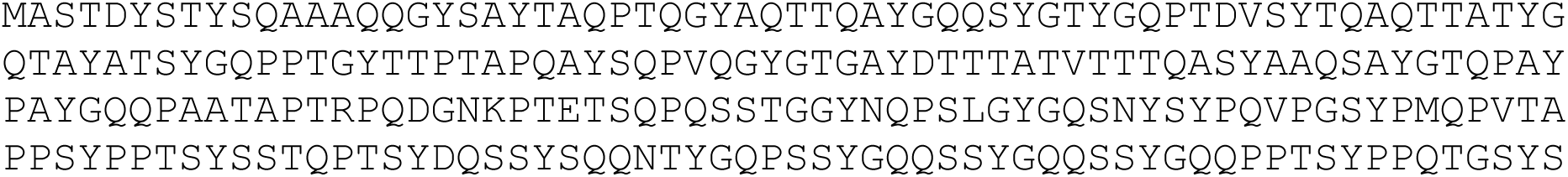
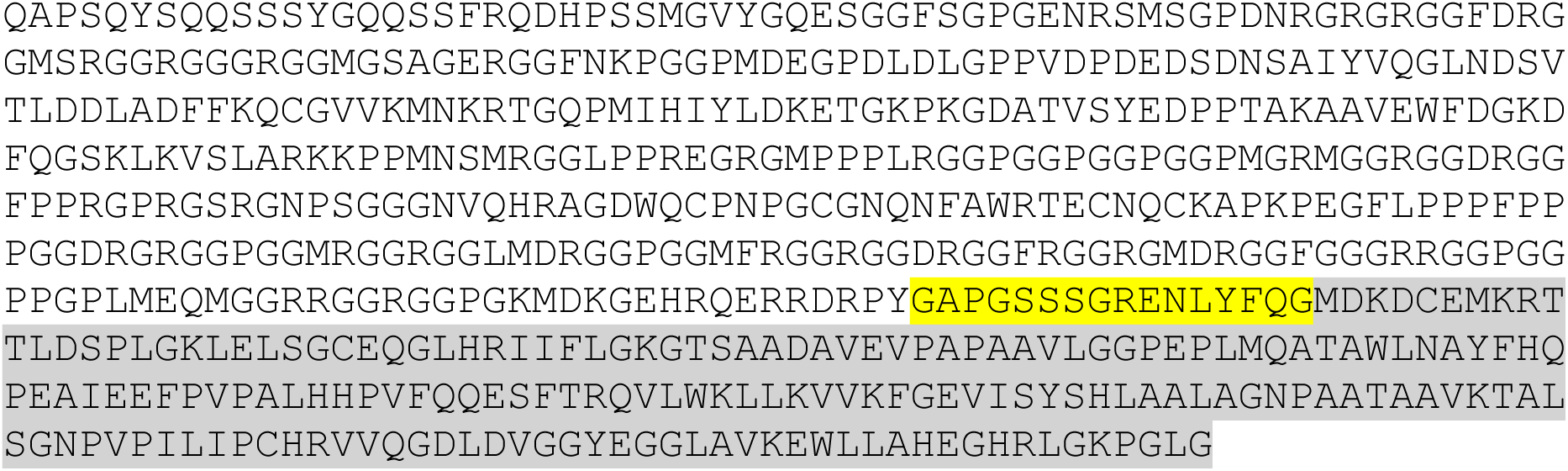

### Methods

#### Protein Purification

Protein purification was carried out following established protocols ^1, 2^. Recombinant proteins were expressed in SF9 insect cells maintained in suspension at 27 °C using ESF921 serum-free medium. The FlexiBAC system facilitated baculovirus production, enabling target gene expression under the control of the polyhedrin promoter. In a typical experiment, 1 L of SF9 cells at a density of 1 million cells per mL were infected with 5 mL of P2 virus and harvested 72 hours post-infection. Cells were collected by centrifugation at 300 RCF for 15 minutes at 4 °C, and the pellet was resuspended in 30 mL of ice-cold lysis buffer (50 mM HEPES pH 7.4, 1 M KCl, and 5% glycerol) supplemented with EDTA-free protease inhibitor tablets. Cell disruption was achieved by sonication on ice for 5 minutes at 35% output with a 50% duty cycle. Cellular debris was removed by centrifugation at 39,800 RCF for 30 minutes at 4 °C.

Subsequent purification steps were performed at room temperature. The cleared lysate was passed through a 5 mL Ni-NTA agarose column (Protino, Macherey-Nagel) using a peristaltic pump. The column was washed with 10 column volumes of lysis buffer containing 10 mM imidazole. Bound protein was eluted with 5 column volumes of elution buffer (50 mM HEPES pH 7.4, 1 M KCl, 5% glycerol, and 300 mM imidazole). Peak fractions were incubated with 10 mL of amylose resin for 10 minutes, followed by gravity flow drainage. The resin was washed with 10-column volumes of lysis buffer, and the target protein was eluted using 5-column volumes of maltose-containing buffer (50 mM HEPES pH 7.4, 1 M KCl, 5% glycerol, and 30 mM maltose). Protein concentration was monitored using a Bradford assay, and purity was assessed by SDS-PAGE.

To remove the N-terminal His-MBP tag, 3C precision protease was added at a 1:100 molar ratio, and digestion was carried out at room temperature for 4 hours. The sample was further purified using size-exclusion chromatography on an ÄKTA system (GE Healthcare) with a Superdex 200 10/300 Increase column pre-equilibrated with storage buffer (50 mM HEPES pH 7.4, 500 mM KCl, 5% glycerol, and 1 mM DTT). Peak fractions of C-terminal SNAP-tagged protein were collected, concentrated, and prepared for experiments.

For the preparation of untagged protein, TEV protease was added at a 1:50 molar ratio to prepare untagged protein and incubated at room temperature for 6 hours. Gel filtration chromatography purified the resulting protein using a Superdex 200 10/300 Increase column equilibrated with storage buffer. Peak fractions were pooled and concentrated using a 30 kDa molecular weight cut-off (MWCO) filter at 3,000 RCF at room temperature. Protein concentration was determined using a NanoDrop ND-1000 spectrophotometer (Thermo Scientific), with 260/280 ratios ranging from 0.52 to 0.56. Peak fractions were pooled and immediately used for dynamic light scattering (DLS) or nanoparticle tracking analysis (NTA) experiments.

SNAP-tagged and untagged proteins were snap-frozen in liquid nitrogen and stored at −80°C.

#### Preparation of ATP.Mg^2+^

Following the previous report ^3^, nucleotide-magnesium complexes were prepared. Briefly, molar equivalent magnesium acetate was added to the solutions of adenosine triphosphate sodium salt and adenosine diphosphate sodium salt and incubated for 5 minutes before the experiments.

#### DLS Measurements

DLS measurements were carried out at 24 °C using a Wyatt DynaPro® Nanostar (Waters|Wyatt Technology, US) instrument. Disposable cuvettes (Uvette 220-1600 nm) from Eppendorf were employed, with a sample volume of 100 µl. A 658 nm laser illuminated the sample solutions, and the intensity of light scattered at an angle of 90° was measured using a single photon counting module. This allows for the measurement of time-dependent fluctuations in the intensity of scattered light as scatterers undergo Brownian motion. The rate of fluctuations is directly related to the diffusion rate of the molecule through the solvent, which in turn correlates with the particles’ hydrodynamic radii. Analyzing intensity fluctuations enables the determination of the diffusion coefficients (D_t_) of particles, which are converted into a size distribution using the Stokes-Einstein equation (equation 1).

Stokes-Einstein relation:

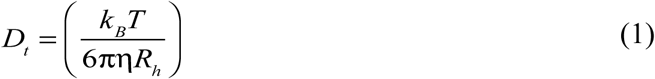

Here, *k_B_* is the Boltzmann constant (1.381 × 10^−23^J/K) and ρι is the absolute (or dynamic) viscosity of the solvent. In this work, we used the hydrodynamic diameter *d_h_* (i.e., *d_h_*= 2*R_h_*) as the preferred way to quantify particle sizes.

The analyte’s translational diffusion coefficient (D_t_) is obtained by automated nonlinear least squares fitting of the autocorrelation function that quantitatively describes the measured time-dependent fluctuations in light scattering intensity. The analysis is done directly using the accompanying DYNAMICS™ software.

For the measurements, all buffer solutions were filtered using 0.2 μm membranes (Millex®-GS units) purchased from Millipore™. All protein samples were centrifuged at 10,000 RCF before measurements. All experiments were conducted with the following settings: Material – protein; Dispersant – buffers; Temperature: 24 °C with equilibration time – 120 seconds, Measurement angle: 90°. Each spectrum represents the average of 12 scans, each of 10 seconds in duration. The samples were prepared by adding centrifuged stock proteins followed by dilution buffer and mixed thoroughly by pipetting three to four times. The samples were equilibrated for 2 minutes at 24 °C, and the data were recorded in 2-minute intervals. DYNAMICS™ (Waters|Wyatt Technology, USA) software is used to control experimental parameters, collect data, and analyze them. The autocorrelation, intensity size distributions, and mean hydrodynamic diameters of each measurement were exported from DYNAMICS™ software and plotted in GraphPad Prism software.

In this report, we present only the intensity size distribution, as it directly correlates with the autocorrelation function without requiring any data transformation as FET proteins mesoscale clusters are polydisperse and quasi-spherical ^1^. For volume transformation, assuming spherical particles, the particle volume is proportional to the cube of the size. For number transformation, assuming isotropic small particles, the scattering intensity from a spherical particle is proportional to the sixth power of the size.

#### Nanoparticle Tracking Analysis (NTA)

Nanoparticle tracking analysis was conducted using the NanoSight Pro from Malvern Instruments, which has a measurement range of 10 nm to 1 μm. The system included a NanoSight syringe pump to inject samples for the experiments. NTA measurements leverage the properties of light scattering and Brownian motion to determine the size distributions and concentrations of particles in liquid suspension. A laser beam (488 nm) was directed through the sample chamber, and the suspended particles were visualized using a 20x magnification microscope. A video file capturing the movement of particles under Brownian motion was recorded using a camera mounted on the microscope, operating at 30 frames per second. The software tracks the movement of individual particles from frame to frame in order to calculate the mean square displacement. The Stokes-Einstein equation (Equation 1) is applied to calculate the particles’ hydrodynamic diameter. The instrument provided the particle concentration by counting particles per frame in known observation volume.

All buffers were filtered through a 0.22 μm polyvinylidene fluoride membrane filter (Merck, Germany). Protein stock solutions were centrifuged at 20000 RCF for 5 minutes at room temperature, and the concentration was measured prior to further measurements. Samples were prepared by adding the centrifuged stock proteins followed by a dilution buffer and mixed thoroughly by pipetting 4 to 6 times. The samples were equilibrated for 2 minutes at 24 °C, and data were recorded approximately 6 minutes after sample preparation.

#### Calculation of Volume Fraction

Nanoparticle Tracking Analysis (NTA) provides particles’ hydrodynamic diameters along with their concentration in particles per milliliter (particles/mL).

The volume fraction of clusters (ϕ) is calculated as: 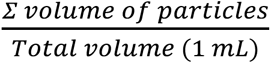

#### Nano Differential Scanning Fluorimetry (nanoDSF)

The thermal unfolding of FUS-SNAP was carried out using nanoDSF with a Prometheus NT.48 (NanoTemper Technologies, Munich, Germany) instrument. Protein samples were prepared in buffers and briefly centrifuged (5 min, 10,000 RCF) to eliminate protein aggregates. Samples were loaded into high-sensitivity glass capillaries (Cat#PR-C006, NanoTemper Technologies, Munich, Germany) and subjected to a linear thermal ramp from 20 °C to 95 °C at a rate of 1 °C/min. Intrinsic protein fluorescence emission was collected at 330 and 350 nm with a dual-UV detector over a temperature gradient. The fluorescence intensity ratio (350/330) was plotted against temperature, and the inflection point of the transition was obtained from the maximum of the first derivative for each measurement using Therm-Control Software (NanoTemper Technologies, Munich, Germany). All experiments were conducted in triplicate; the mean and standard deviation were calculated for all three measurements.

**Supporting Table 2:** Comparison of hydrodynamic diameter (d*ₕ*) measurements obtained from dynamic light scattering (DLS) and nanoparticle tracking analysis (NTA). DLS data includes the polydispersity index (PDI) of the mesoscale clusters, and NTA data includes the volume fractions of the clusters.

|  | DLS |  |  |  | NTA |  |
| --- | --- | --- | --- | --- | --- | --- |
| | Mean size<br>(nm)<br>@ 30 mins | PDI | Mean size<br>(nm)<br>@ 8 mins | PDI | Mean size<br>(nm)<br>@ ~8 mins | Volume<br>fractions<br>($\phi_{\text{cluster}} \times 10^{-5}$) |
| 0.25 $\mu\text{M}$<br>FUS SNAP | 281.74<br>$\pm 14.98$ | 0.122<br>$\pm 0.048$ | 244.32<br>$\pm 11.98$ | 0.132<br>$\pm 0.052$ | 225<br>$\pm 14.2$ | 0.209<br>$\pm 0.033$ |
| 0.5 $\mu\text{M}$<br>FUS SNAP | 441.80<br>$\pm 74.15$ | 0.119<br>$\pm 0.036$ | 344.41<br>$\pm 35.88$ | 0.125<br>$\pm 0.061$ | 282<br>$\pm 20.5$ | 0.448<br>$\pm 0.145$ |
| 1 $\mu\text{M}$<br>FUS SNAP | 1,012.40<br>$\pm 72.95$ | 0.172<br>$\pm 0.052$ | 705.29<br>$\pm 107.69$ | 0.157<br>$\pm 0.099$ | 493<br>$\pm 17.32$ | 2.407<br>$\pm 0.835$ |
| 0.25 $\mu\text{M}$<br>FUS SNAP<br>with 1 mM<br>ATP.Mg<sup>2+</sup> | 673.45<br>$\pm 104.06$ | 0.167<br>$\pm 0.065$ | 476.28<br>$\pm 31.73$ | 0.114<br>$\pm 0.022$ | 248.5<br>$\pm 10.96$ | 0.305<br>$\pm 0.116$ |
| 0.25 $\mu\text{M}$<br>FUS SNAP<br>with 2 mM<br>ATP.Mg<sup>2+</sup> | 642.58<br>$\pm 70$ | 0.127<br>$\pm 0.054$ | 446.10<br>$\pm 30.72$ | 0.137<br>$\pm 0.056$ | 267.3<br>$\pm 16.19$ | 0.392<br>$\pm 0.279$ |
| 0.25 $\mu\text{M}$<br>FUS SNAP<br>with 5 mM<br>ATP.Mg<sup>2+</sup> | 312.38<br>$\pm 45.93$ | 0.146<br>$\pm 0.059$ | 323.25<br>$\pm 60.26$ | 0.153<br>$\pm 0.053$ | 174<br>$\pm 12.85$ | 0.126<br>$\pm 0.062$ |
| 0.25 $\mu\text{M}$<br>FUS SNAP<br>with 10 mM<br>ATP.Mg<sup>2+</sup> | 222.21<br>$\pm 69.29$ | 0.318<br>$\pm 0.212$ | 169.67<br>$\pm 30.62$ | 0.279<br>$\pm 0.19$ | 155.5<br>$\pm 11.84$ | 0.0568<br>$\pm 0.044$ |

**Supporting Figure 1:**
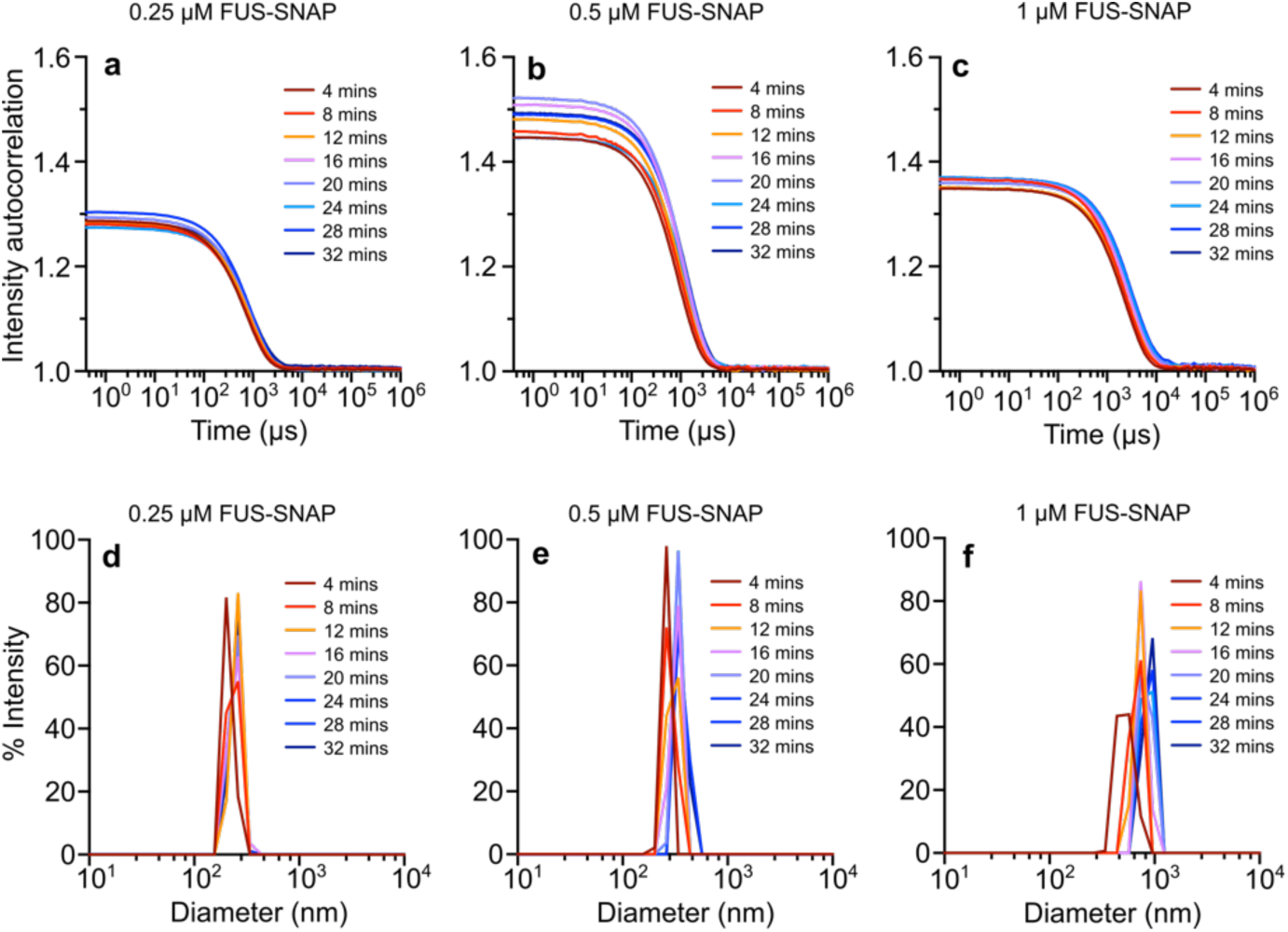
The autocorrelation function from DLS of solutions with varying concentrations of FUS-SNAP: 0.25 μM (a), 0.5 μM (b), and 1 μM (c) in 20 mM HEPES, pH 7.4, with 10 mM KCl. The size distributions of the scatterers are presented as % intensity derived from the corresponding intensity autocorrelation of DLS samples containing 0.25 μM (d), 0.5 μM (e), and 1 μM (f) FUS-SNAP.

**Supporting Figure 2:**
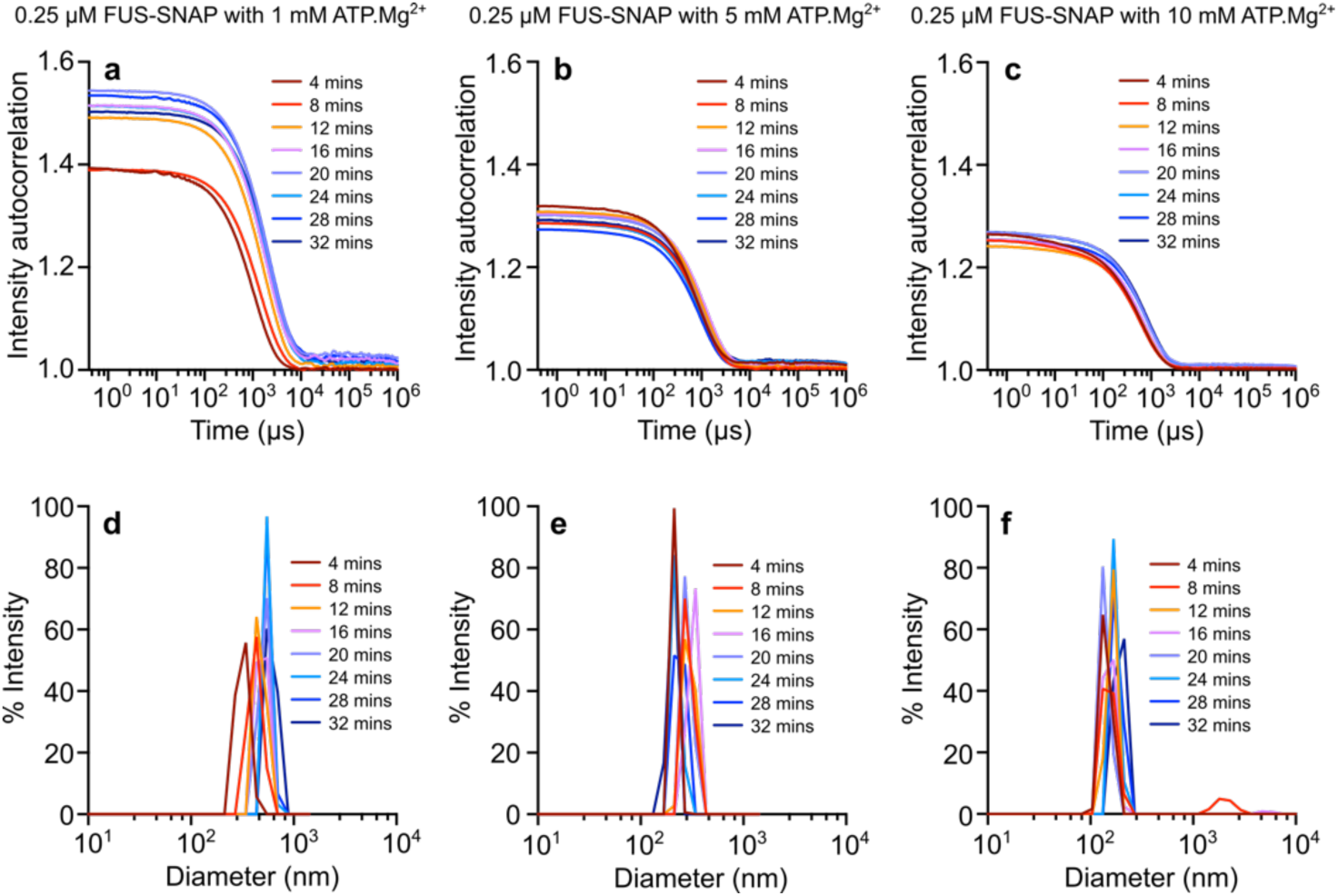
The autocorrelation function from DLS of solutions containing 0.25 μM FUS-SNAP with different concentrations of ATP.Mg^2+^, 1 mM (a), 5 mM (b), and 10 mM (c) in 20 mM HEPES, pH 7.4, with 10 mM KCl. The size distributions of the scatterers are shown as percentage intensity derived from the intensity autocorrelation of DLS samples containing 1 mM (d), 5 mM (e), and 10 mM (f) ATP.Mg^2+^ with 0.25 μM FUS-SNAP.

**Supporting Figure 3:**
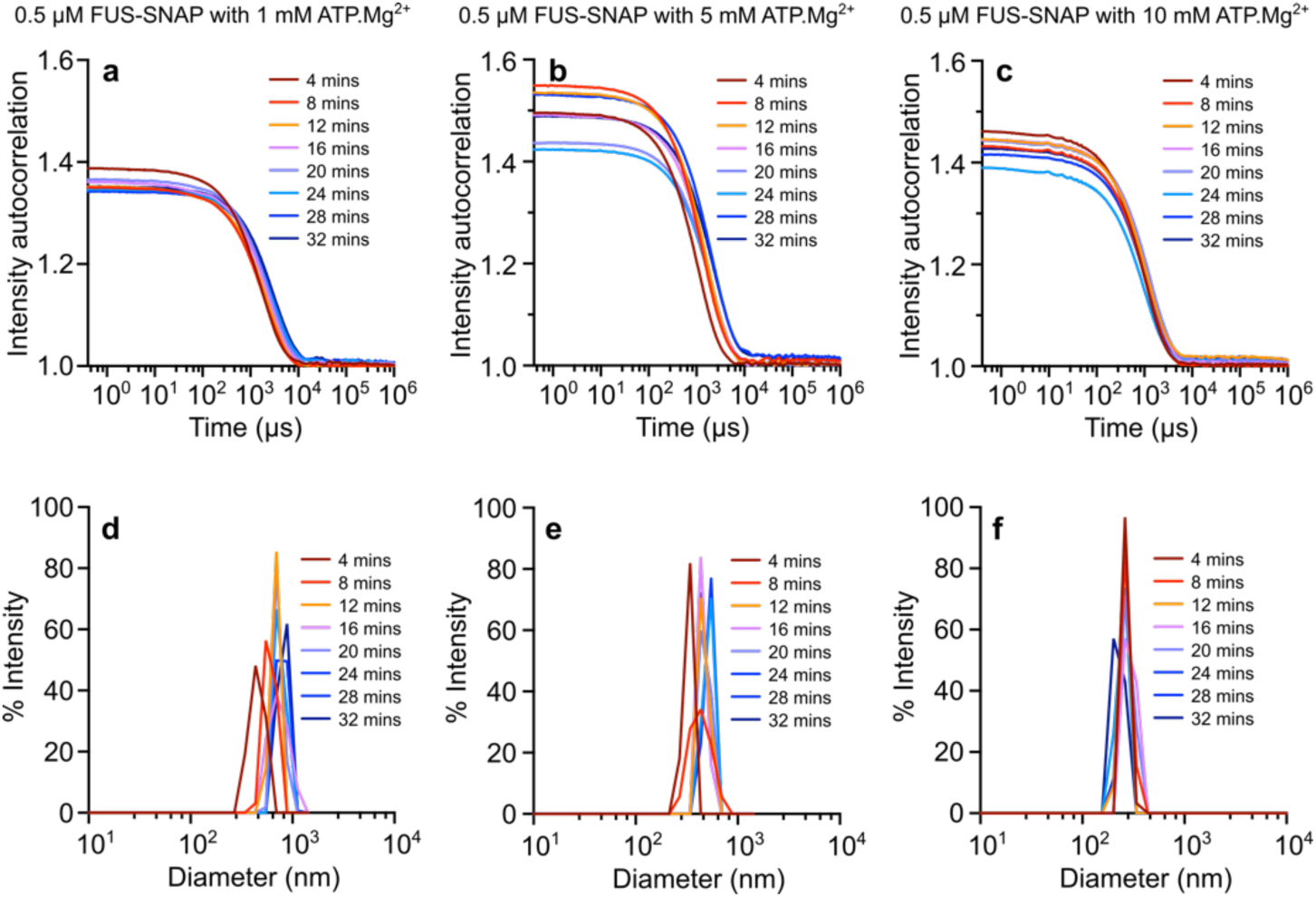
The autocorrelation function from DLS of solutions containing 0.5 μM FUS-SNAP with varying concentrations of ATP.Mg^2+^, 1 mM (a), 5 mM (b), and 10 mM (c), in 20 mM HEPES, pH 7.4, with 10 mM KCl. The size distributions of the scatterers are presented as % intensity derived from the intensity autocorrelation of DLS samples containing 1 mM (d), 5 mM (e), and 10 mM (f) ATP.Mg^2+^ with 0.5 μM FUS-SNAP.

**Supporting Figure 4:**
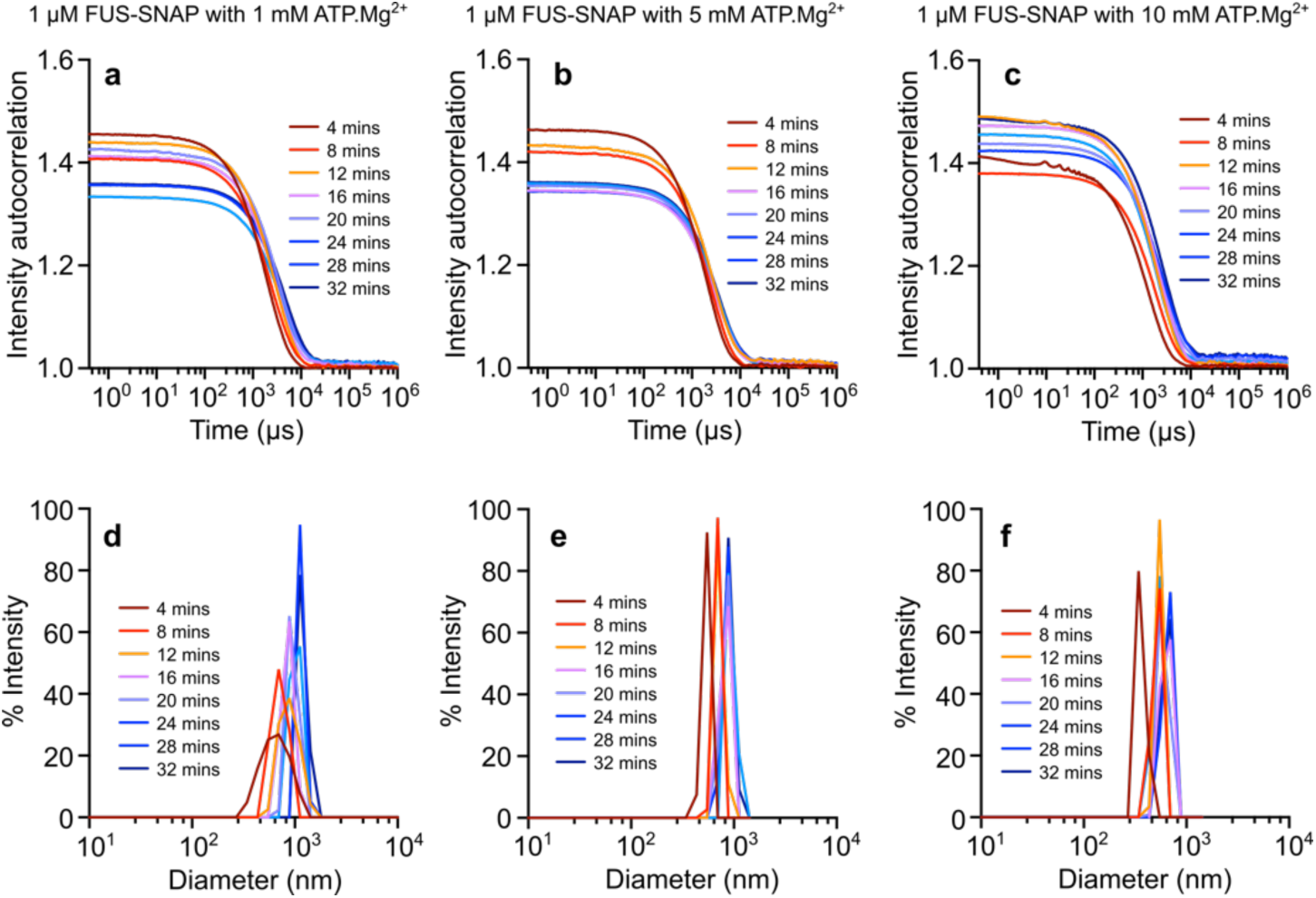
The autocorrelation function from DLS of solutions containing 1 μM FUS-SNAP with varying concentrations of ATP.Mg^2+^: 1 mM (a), 5 mM (b), and 10 mM (c) in 20 mM HEPES, pH 7.4, with 10 mM KCl. The size distributions of the scatterers are shown as % intensity derived from the intensity autocorrelation of DLS samples containing 1 mM (d), 5 mM (e), and 10 mM (f) ATP.Mg^2+^ with 1 μM FUS-SNAP.

**Supporting Figure 5:**
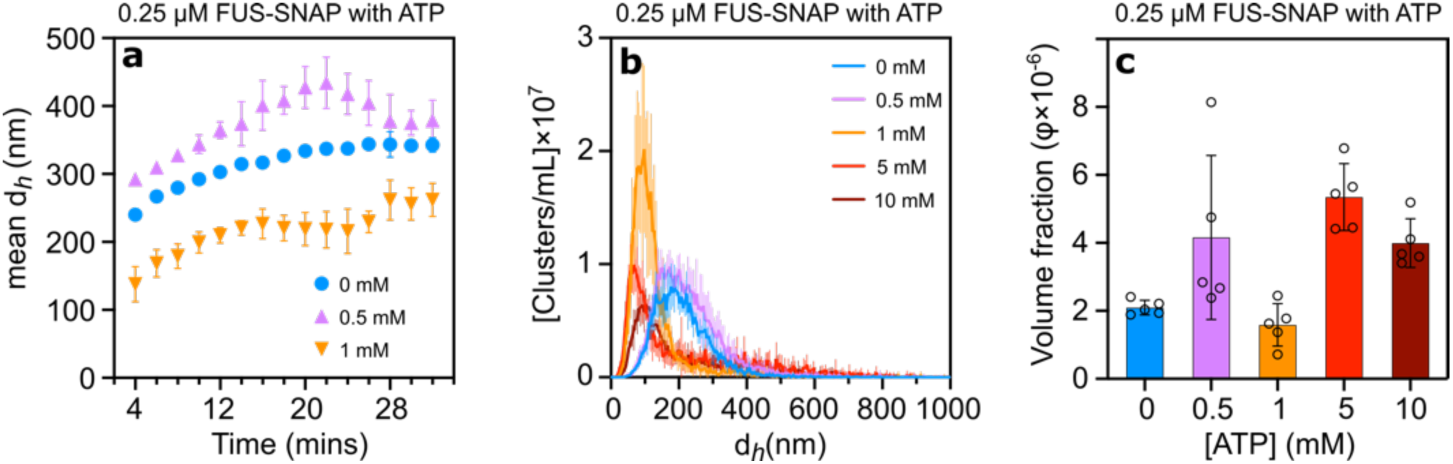
Adenosine triphosphate (ATP) without Mg^2+^ ion regulates the cluster size of sub-saturation FUS-SNAP clusters. Dynamic Light Scattering data show the hydrodynamic diameter (d_h_) of mesoscale clusters at 1 µM FUS-SNAP over 32 minutes with various concentrations of ATP (a). NTA data for cluster hydrodynamic diameter at 0.25 μM FUS-SNAP with different ATP concentrations are shown in (b), while (c) displays the corresponding relative abundance of clusters as volume fraction. Data are presented as mean ± SD, with n=3 (DLS) and n=5 (NTA) independent samples.

**Supporting Figure 6:**
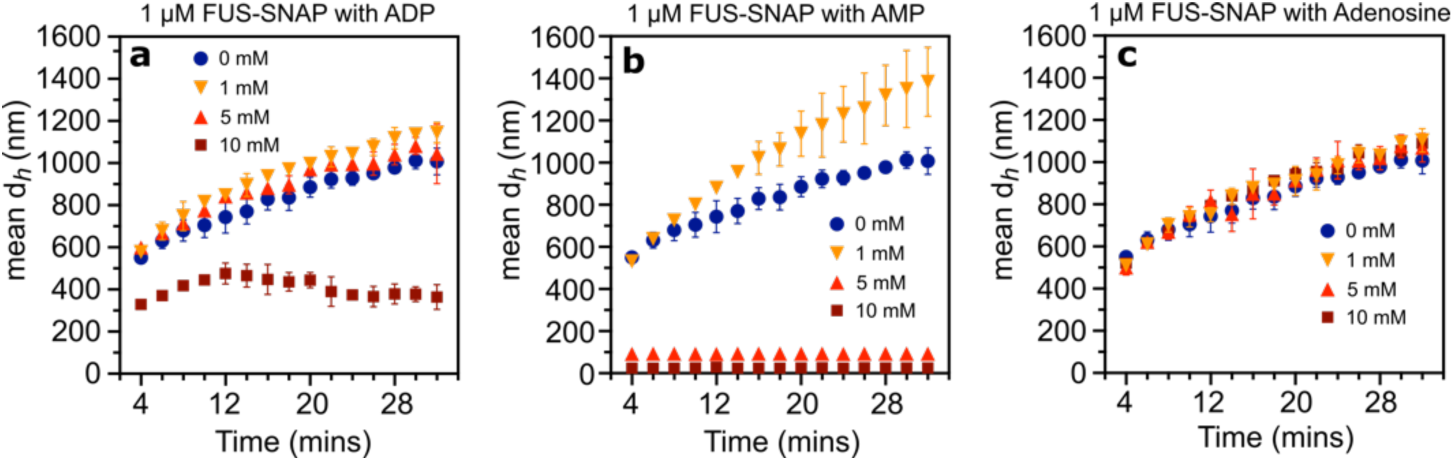
Adenosine diphosphate (ADP) and adenosine monophosphate (AMP) regulate the cluster size of sub-saturation FUS-SNAP clusters, while adenosine does not. Dynamic Light Scattering data show the hydrodynamic diameter (d_h_) of mesoscale clusters at 1 µM FUS-SNAP over 32 minutes with various concentrations of ADP (a), AMP (b), and adenosine (c). Three independent samples (n=3) were used for the measurements, and the data are presented as mean values ± SD.

**Supporting Figure 7:**
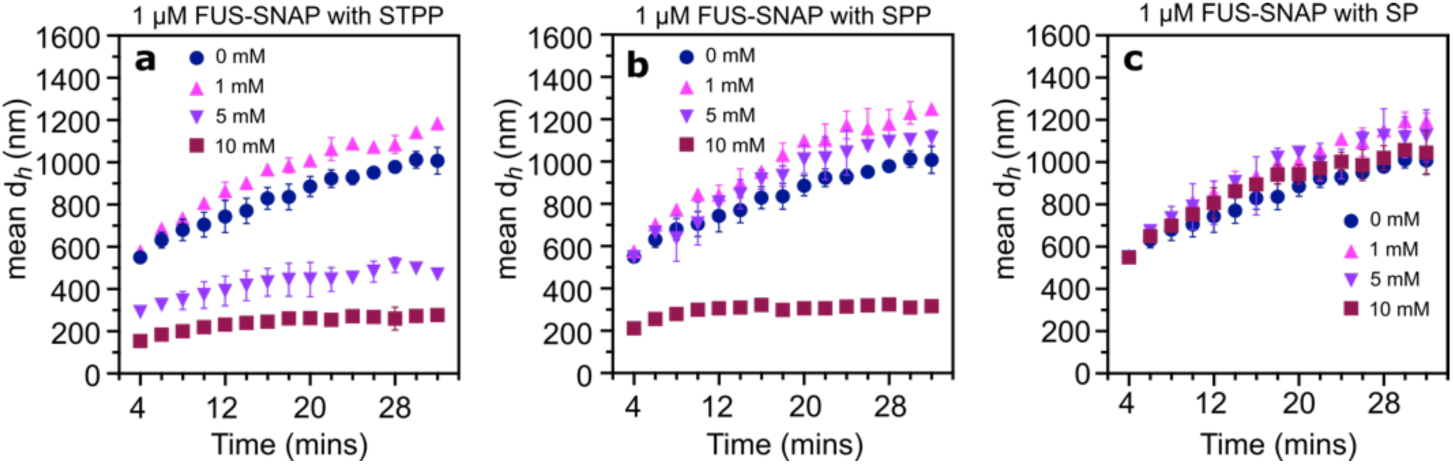
Phosphates are crucial in regulating the size of sub-saturated FUS-SNAP clusters. Dynamic Light Scattering data show the hydrodynamic diameter (*d_h_*) of mesoscale clusters at 1 µM FUS-SNAP over 32 minutes with varying concentrations of sodium tripolyphosphate (STPP) (a), sodium pyrophosphate (SPP) (b), and sodium phosphate (SP) (c). Three independent samples (n=3) were used for the measurements, and the data are presented as mean values ± SD.

**Supporting Figure 8:**
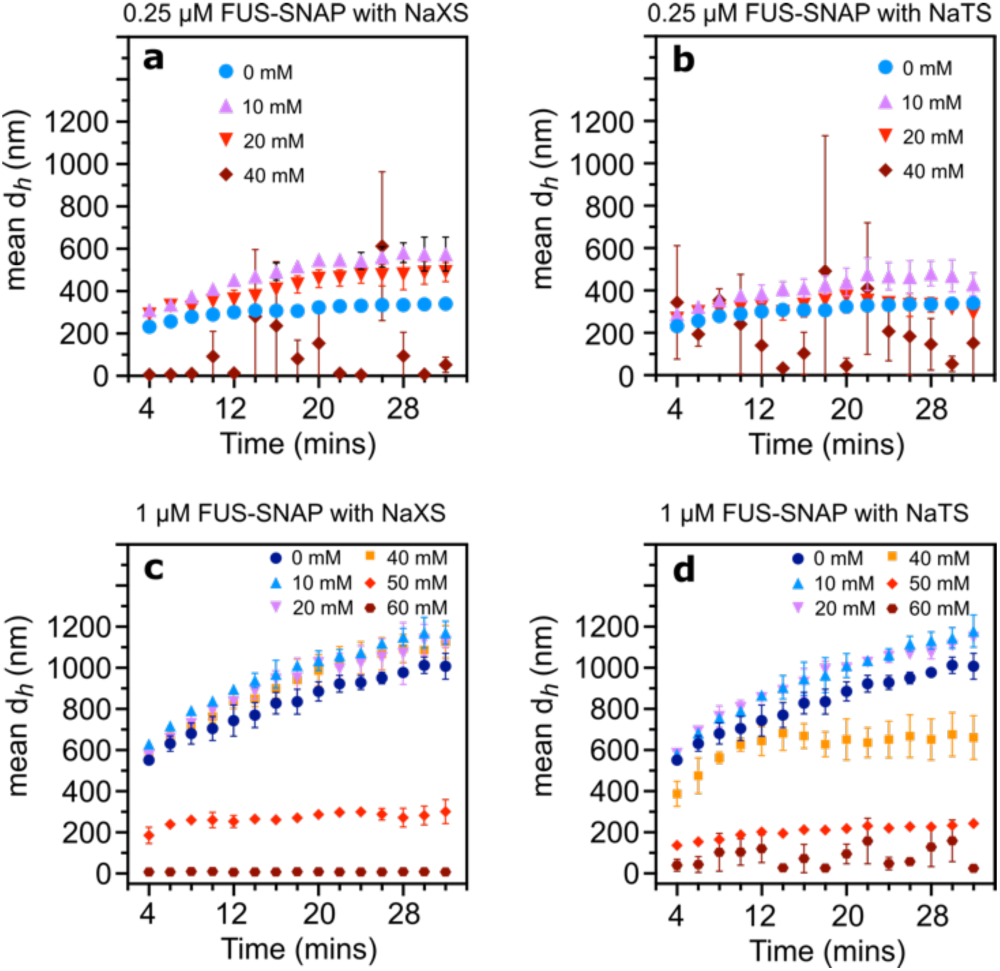
Hydrotropes modulate the size of saturation clusters. Dynamic light scattering data show the hydrodynamic diameter (*d_h_*) of mesoscale clusters of 0.25 µM FUS-SNAP over 32 minutes with various concentrations of Sodium Xylene Sulfate (NaXS) (a) and Sodium Toluene Sulfate (NaTS) (b). Dynamic light scattering data show the hydrodynamic diameter (*d_h_*) of mesoscale clusters at 1 µM FUS-SNAP over 32 minutes with various concentrations of Sodium Xylene Sulfate (NaXS) (c) and Sodium Toluene Sulfate (NaTS) (d). Three independent samples (n=3) were used for the measurements, and the data are presented as mean values ± SD.

**Supporting Figure 9:**
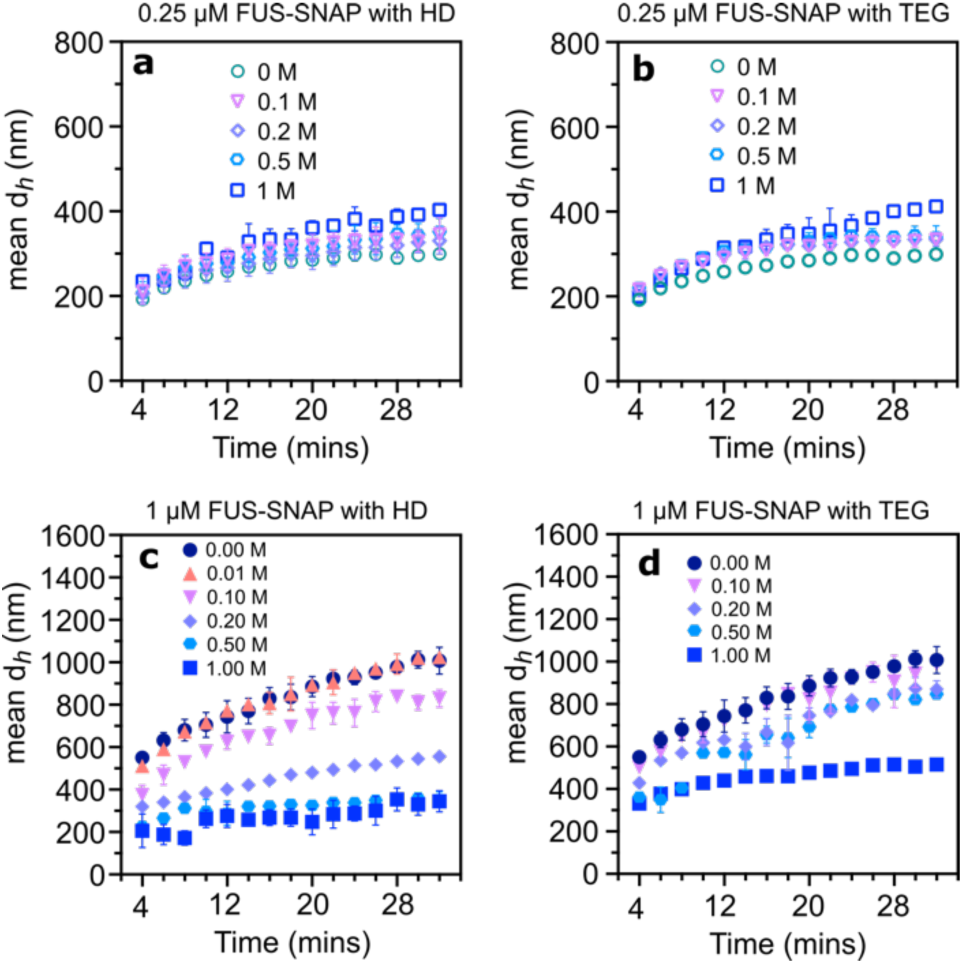
Hexanediol (HD) and triethylene glycol (TEG) influence the size of condensates differently at different concentrations of FUS-SNAP. Dynamic light scattering data show the hydrodynamic diameter (*d_h_*) of mesoscale clusters of 0.25 µM FUS-SNAP over 32 minutes with different concentrations of HD (a) and TEG (b). Additionally, dynamic light scattering data indicate the hydrodynamic diameter (*d_h_*) of mesoscale clusters at 1 µM FUS-SNAP over 32 minutes with varying concentrations of HD (c) and TEG (d). Three independent samples (n=3) were used for the measurements, and the data are presented as mean values ± SD.

**Supporting Figure 10:**
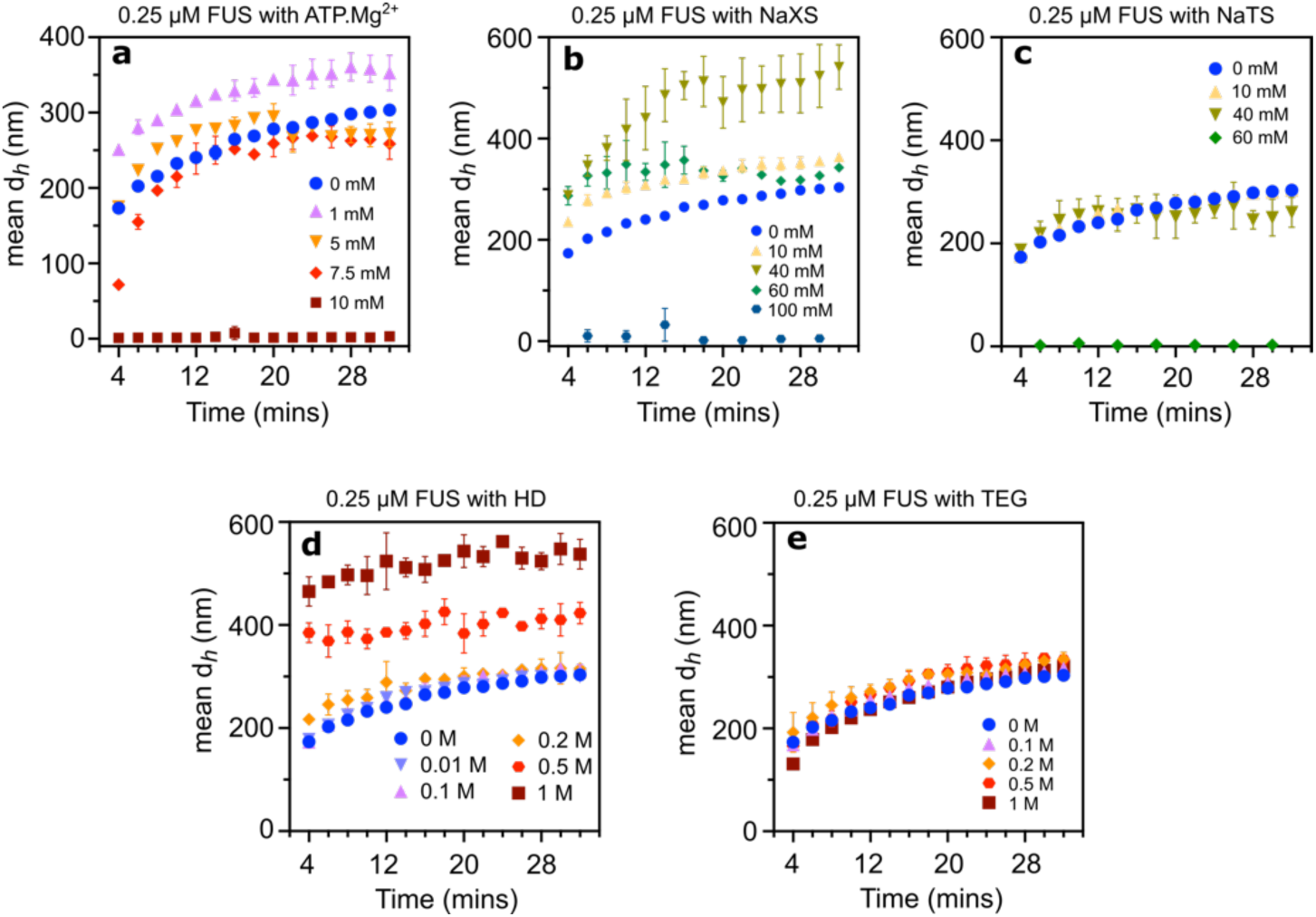
Sub-saturation cluster formation of untagged FUS is similar to FUS-SNAP in the presence of ATP and hydrotropes but differs with HD and TEG. Dynamic Light Scattering data show mesoscale clusters’ hydrodynamic diameter (d_h_) at 0.25 μM FUS over 32 minutes with various concentrations of ATP.Mg^2+^ (a), NaXS (b), NaTS (c), HD (d), and TEG (e). Three independent samples (n=3) were used for the measurements, and the data are presented as mean values ± SD.

**Supporting Figure 11:**
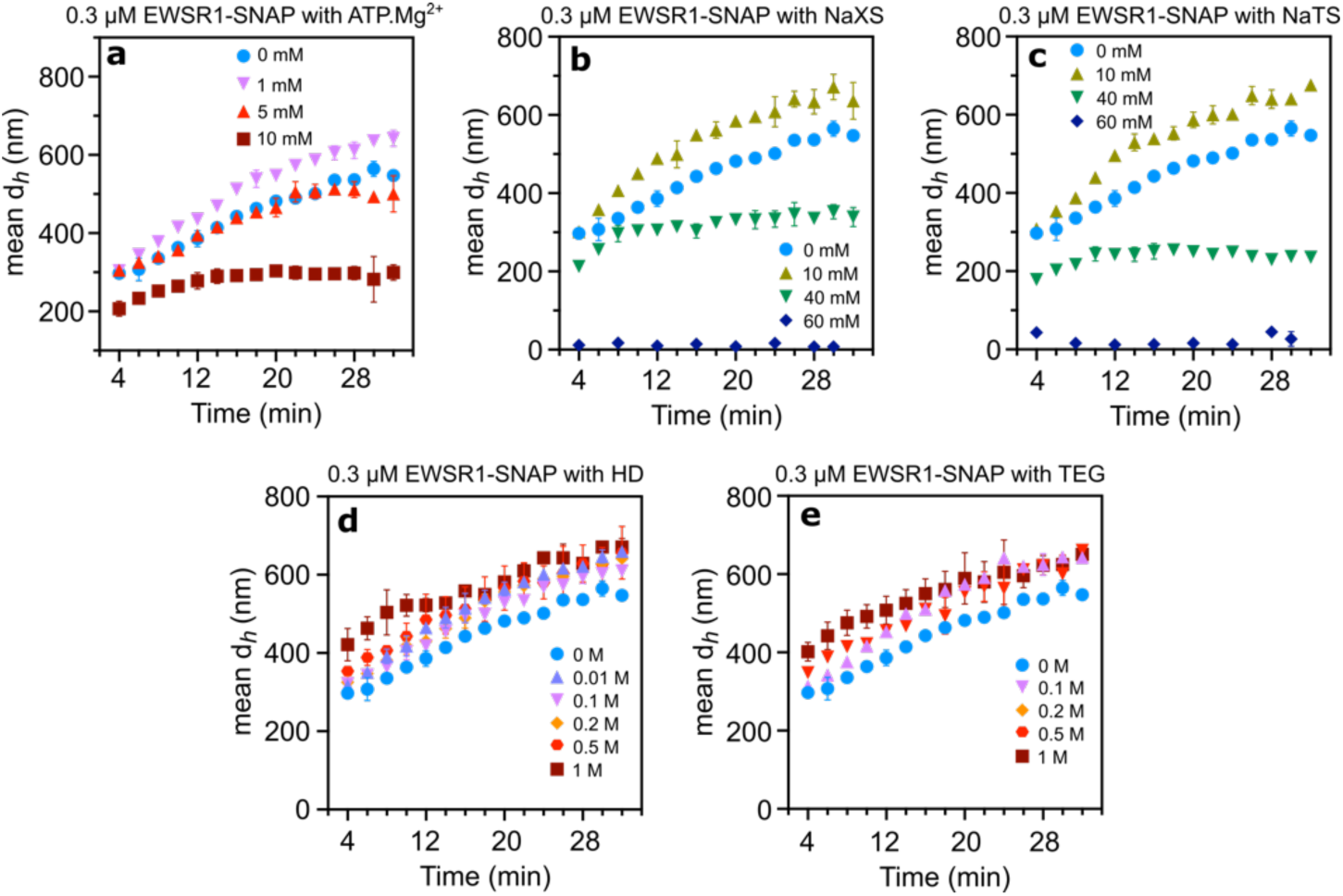
Sub-saturation cluster formation of EWSR1-SNAP is also influenced by the presence of ATP and hydrotropes but differs with HD and TEG. Dynamic Light Scattering data show mesoscale clusters’ hydrodynamic diameter (d_h_) at 0.3 μM EWSR1-SNAP over 32 minutes with various concentrations of ATP.Mg^2+^ (a), NaXS (b), NaTS (c), HD (d), and TEG (e). Three independent samples (n=3) were used for the measurements, and the data are presented as mean values ± SD.

**Supporting Figure 12:**
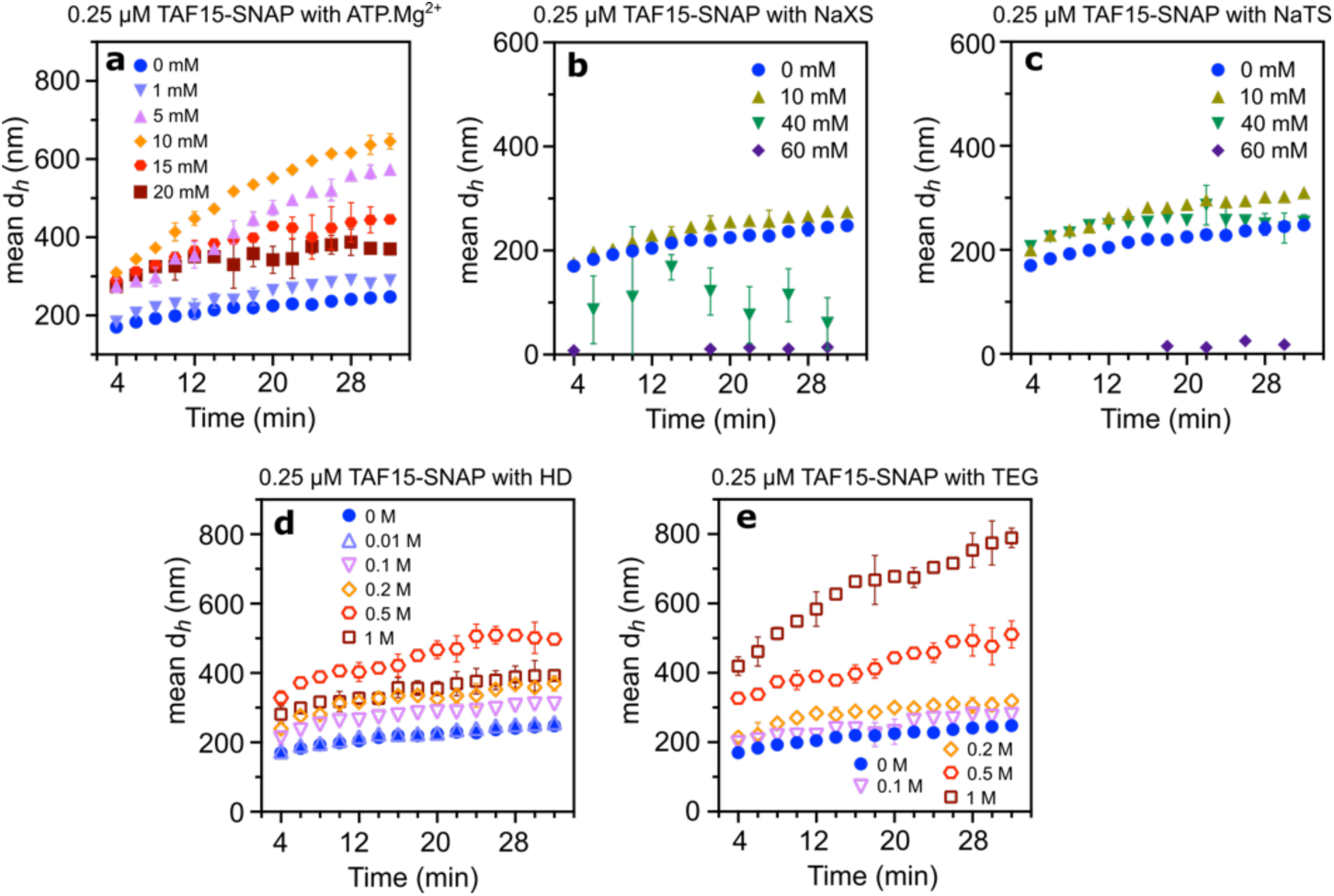
Sub-saturation cluster formation of TAF15-SNAP is also influenced by the presence of ATP and hydrotropes but differs with HD and TEG. Dynamic Light Scattering data show mesoscale clusters’ hydrodynamic diameter (d_h_) at 0.25 μM TAF15-SNAP over 32 minutes with various concentrations of ATP.Mg^2+^ (a), NaXS (b), NaTS (c), HD (d), and TEG (e). Three independent samples (n=3) were used for the measurements, and the data are presented as mean values ± SD.

**Supporting Figure 13:**
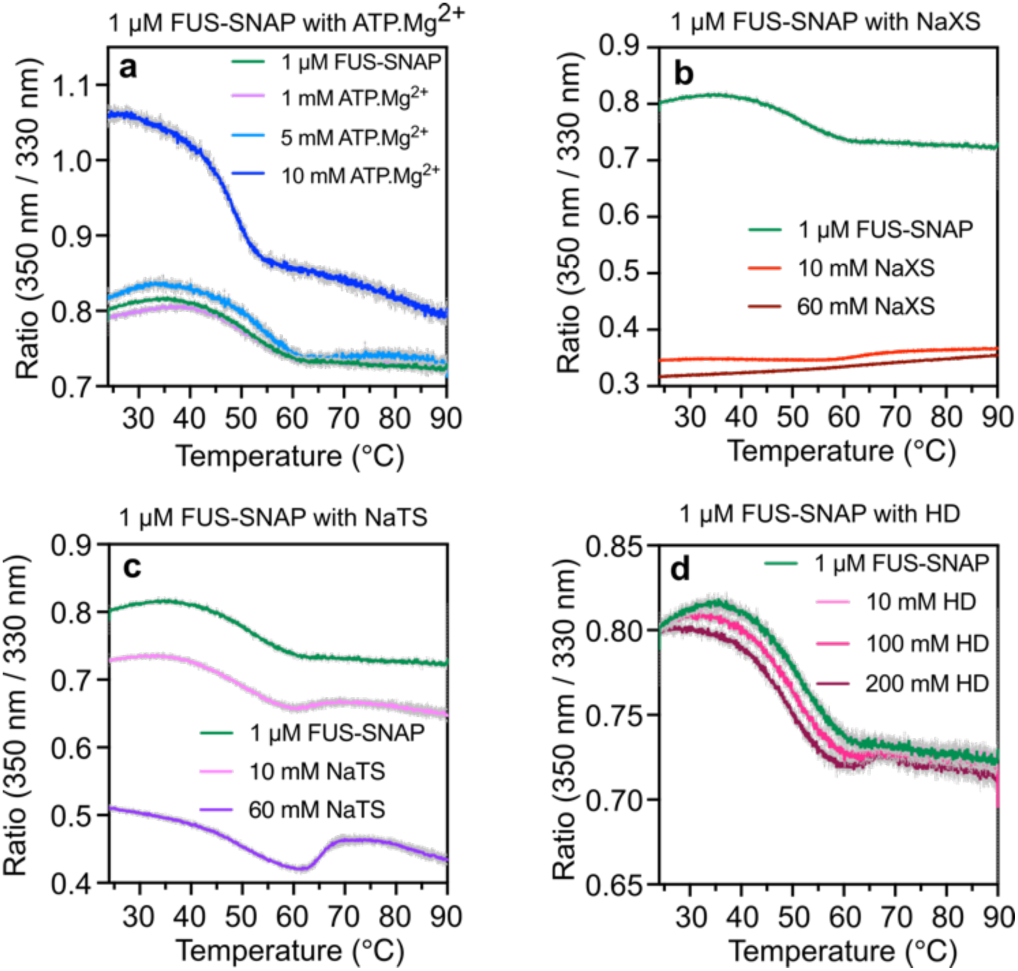
Interactions with ATP and other small amphiphilic molecules alter the transition temperature of FUS-SNAP probed by NanoDSF. The unfolding curves (350/330) ratio with the apparent transition temperatures of FUS-SNAP with ATP.Mg^2+^ (a), NaXS (b), NaTS (c), and HD (d).

